# Prevalence and Actionability of MTAP Loss in Oncogene-Driven Lung Cancer

**DOI:** 10.64898/2026.01.21.700721

**Authors:** Takashi Sakai, Zofia Piotrowska, Charlotte I. Wang, Beow Y. Yeap, Rebecca S. Heist, Jessica J. Lin, Lauren E. Highfield, Jennifer L. Petersen, Mandeep Banwait, Joyce Liang, Manisha Madhavan, Aaron N. Hata, Mari Mino-Kenudson, Ibiayi Dagogo-Jack

**Author notes:** Equal contribution (Aaron N. Hata, Mari Mino-Kenudson, Ibiayi Dagogo-Jack). **Corresponding Author:** Ibiayi Dagogo-Jack, MD, Massachusetts General Hospital, Department of Medicine, 55 Fruit Street, Boston, MA, USA, 02114.

## Abstract

**Background:** Methylthioadenosine phosphorylase (MTAP) loss co-occurs with actionable genomic alterations in non-small cell lung cancer (NSCLC) and creates vulnerability to protein arginine methyltransferase 5 (PRMT5) inhibition. Estimates of prevalence of MTAP loss rely on next-generation sequencing which can underestimate copy losses. Moreover, the activity of PRMT5 inhibitors in oncogene-driven NSCLC is not well established.

**Methods:** We assessed MTAP expression by immunohistochemistry (IHC) in 243 NSCLCs (n=132 early stage, n=111 metastatic), including 33 specimens with paired lymph nodes. Antiproliferative activity of PRMT5 inhibitor monotherapy and in combination with tyrosine kinase inhibitors (TKIs) was evaluated in MTAP-deleted NSCLC cell lines harboring *EGFR* or *KRAS* mutations or *ALK* rearrangements.

**Results:** Among 243 NSCLC specimens from 240 patients (72% with driver alterations, 90% adenocarcinoma), MTAP loss was identified in 43 (18%) specimens from 40 (17%) patients, including 18 (14%) early-stage and 22 (20%) metastatic tumors. MTAP loss occurred in 24% of stage 4 driver-positive NSCLCs versus 14% of driver-negative tumors (p=0.314). Twenty (61%) lung-nodal pairs demonstrated concordance; eight cases only exhibited decreased MTAP expression in nodes. Variable sensitivity to PRMT5 inhibitors was observed in 22 MTAP-deleted NSCLC cell lines (9 *EGFR*-mutant, 7 *KRAS*-mutant, 6 *ALK*-rearranged), with responses seen in TKI-sensitive and TKI-resistant lines. SDMA (symmetric dimethylarginine) expression did not predict PRMT5 inhibitor sensitivity. TKI + PRMT5 inhibitor combos had greater activity than monotherapy.

**Conclusions:** MTAP loss occurs in 1-in-5 oncogene-driven metastatic NSCLCs. PRMT5 inhibitor activity is independent of TKI exposure, driver alteration, and SDMA expression and enhanced by addition of TKI. These findings support clinical evaluation of PRMT5 inhibitor + TKI combinations for advanced NSCLC.

## INTRODUCTION

Approximately 15% of non-small cell lung cancer (NSCLC) harbors homozygous deletion of the methylthioadenosine phosphorylase (*MTAP*) gene.^1–3^ *MTAP* deletions can encompass all eight exons or selectively engulf a subset of exons.^2^ The functional consequence of the diverse permutations of exon deletions depends on which MTAP exons are omitted. For example, focal exon 8 deletion may only partially impair MTAP protein function.^4^ Thus, immunohistochemistry (IHC), which interrogates MTAP protein expression, remains the most direct way to establish loss of MTAP function.^3^ *MTAP* is co-deleted with *CDKN2A* in the majority of NSCLC due to proximity on chromosome 9p21.^1–3^ In addition to *CDKN2A*, *MTAP* deletions occur alongside oncogenic driver alterations in NSCLC, with some studies suggesting that exposure to targeted therapies can promote acquisition of MTAP deletions.^1,2,5^ As prior studies utilized next-generation sequencing (NGS) instead of IHC to determine the prevalence of MTAP loss in oncogene-driven NSCLC, current estimates of the co-existence of MTAP loss and oncogenic alterations may be imprecise. Moreover, as *MTAP* deletion can be a subclonal event, it remains to be determined whether MTAP loss is uniformly present in distinct anatomic sites.

*MTAP* deletion is associated with poor prognosis and decreased sensitivity to molecularly agnostic therapies in pan-tumor analyses.^1^ The impact of MTAP deletion on targeted therapy outcomes in NSCLC has not been fully elucidated. Recent studies suggest decreased durability of osimertinib among patients with *EGFR-*mutant NSCLC with MTAP loss.^5^ *CDKN2A* alterations have also been linked to early progression on ALK, EGFR, and KRAS G12C inhibitors and implicated in resistance to immunotherapy.^6–10^ The adverse predictive and prognostic role of *CDKN2A/MTAP* deletions substantiates efforts to develop selective therapeutic approaches for tumors harboring 9p21 deletions, including synthetic lethal strategies that exploit MTAP’s role in purine salvage. In the purine salvage pathway, MTAP catalyzes metabolism of an intermediate substrate methylthioadenosine (MTA) to re-generate adenine and methionine. MTAP deletion leads to a MTAP deficient, MTA replete state in which the accumulation of MTA partially inhibits the methyltransferase activity of protein arginine methyltransferase 5 (PRMT5), an enzyme involved in epigenetic regulation of proliferation, DNA damage response, and apoptosis.^11^ Impaired function of PRTM5 resulting from MTAP loss imparts a metabolic vulnerability preventing tumors from withstanding further PRMT5 depletion.^12,13^

As proof of concept, monotherapy with investigational MTA-cooperative PRMT5 inhibitors (e.g., AMG193, MRTX1719/BMS-985604) precipitates tumor regressions in patients with MTAP-deleted NSCLC, including osimertinib-resistant, *EGFR-*mutant tumors.^5,14,15^ Given encouraging early findings, ongoing clinical studies are exploring regimens combining PRMT5 inhibitors with established NSCLC targeted therapies. However, the potential of these combinations in tyrosine kinase inhibitor-naïve or resistant oncogene-driven tumors has not been well characterized. Here, we performed IHC to assess MTAP expression in 240 NSCLCs harboring diverse molecular alterations to estimate the prevalence of MTAP loss across molecular contexts, disease sites, and disease stages. We also evaluated the activity of PRMT5 inhibition as monotherapy and in combination with targeted therapies in cell lines harboring *EGFR* mutations, *KRAS* mutations, or *ALK* fusions.

## MATERIALS AND METHODS

### Data Collection and Analysis of MTAP and CDKN2A Status

We retrospectively collected tissue specimens from patients with untreated stage IV lung adenocarcinoma or resected stage I-III lung adenocarcinoma (per AJCC v8.0) who received care at Massachusetts General Hospital between 2021-2024. MTAP expression was determined by IHC using an automated stainer (Bond III; Leica Microsystems, Bannockburn, IL) and the 42-T antibody (1:100, Santa Cruz, sc-100782). MTAP expression was classified as preserved, heterogenous/dichotomous, significantly reduced, and lost based on cytoplasmic staining as determined by a pathologist (M.M.K). To identify *CDKN2A* alterations (mutations and copy number changes) and oncogenic driver alterations, NGS was performed with either the Solid SNAPSHOT assay (DNA NGS) + Solid Fusion Assay (RNA NGS), as previously described (Archer Dx),^16^ or the Genexus Oncomine Precision Assay (OPA; Thermo Fisher).^17^ *MTAP* was not among the genes interrogated in the three assays. For specimens submitted for the Genexus OPA, RNA and DNA were extracted from formalin-fixed paraffin-embedded tumor tissue, with library prepared using an AmpliSeq panel, and analyzed on an Ion Torrent Integrated Sequencer. All non-metastatic NSCLCs were analyzed using the Genexus OPA whereas a combination of approaches was employed for stage IV tumors. Medical records for patients from whom specimens originated were mined to collect demographic information and treatment histories. All patients included in these analyses provided consent for molecular testing. This study was supported by an IRB approved protocol.

### Cell Lines and Cell Culture

All cell lines used in this study are listed in **Supplementary Data Table 3.** The following cell lines— H3255, HCC4006, H1650, H1975, LU99A, SW1573, SKLU1, A549, H2030, H441, SW900, H1944, COR-L23, H1792, H460, H23, A427, H2122, H3122, PC9, HCC827, CALU1, H358, H1573, DV90, H2009, H1155, and LU65 —were obtained from the MGH Center for Molecular Therapeutics. Short tandem repeat (STR) profiling (Biosynthesis, inc.) was performed to verify the identity of each cell line at the time of the study. MGH patient-derived cell lines were established in our laboratory from biopsy or pleural effusion samples, as described.^18^ Informed consent was obtained from all patients under protocols approved by the Dana-Farber/Harvard Cancer Center Institutional Review Board. Most *EGFR*-mutant, *ALK*-positive, and patient-derived cell lines were maintained in RPMI 1640 medium (Thermo Fisher Scientific, USA) supplemented with 10% fetal bovine serum (FBS; Thermo Fisher Scientific, USA). *KRAS*-mutant cell lines were generally cultured in RPMI1640 medium containing 5% FBS. A subset of *KRAS*-mutant lines (SW1573, LU65, A427) was maintained in DMEM/F-12 (1:1) (Thermo Fisher Scientific, USA) supplemented with 5% FBS, while SKLU1 was cultured in DMEM (Gibco, Thermo Fisher Scientific) with 10% FBS. Some *ALK*-positive cell lines, such as MGH979-6.7 and MGH915-3, were maintained in RPMI 1640 medium supplemented with 300 nM alectinib, while MGH073-2 was cultured in RPMI with 300 nM crizotinib. All cells were routinely verified to be free of mycoplasma contamination.

### Drug-Tolerant Persister and Resistant Cells

HCC4006 drug-tolerant persister (DTP) cells were generated by culturing cells in 1 μM osimertinib for 14 days. Cells that remained viable after drug treatment were defined as DTPs. The gefitinib-resistant HCC4006-GR cell line and the lorlatinib-resistant H3122 LR-B cell line were previously established by our group.^18,19^ The RMC-6236-resistant LU99A cell line was generated by culturing cells in medium supplemented with 1 μM RMC-6236 for 6 weeks. Cells exhibiting significantly reduced sensitivity to RMC-6236 in subsequent viability assays were defined as resistant. Resistant cell lines were maintained in 1 μM TKIs and drug were replaced twice per week.

### Antibodies and Reagents

Antibodies used for Western blotting are listed in Supplementary Data Table 2. For cell culture, the following tyrosine kinase inhibitors (TKIs) were used: gefitinib (first-generation EGFR inhibitor; Selleck Chemicals, Houston, TX, USA), osimertinib (third-generation EGFR inhibitor; Selleck Chemicals, Houston, TX, USA), RMC-6236 (pan-KRAS inhibitor; MedChemExpress, Monmouth Junction, NJ, USA), alectinib (second-generation ALK inhibitor; Selleck Chemicals, Houston, TX, USA), crizotinib (first-generation ALK inhibitor; Selleck Chemicals, Houston, TX, USA), and lorlatinib (third-generation ALK inhibitor; MedChemExpress, Monmouth Junction, NJ, USA). All compounds were dissolved in DMSO at a final concentration of 10 mM and stored at −20 °C until use.

### Western Blotting and Quantification

The following primary antibodies were used: anti-CDKN2A (p16^INK4a^) (Cell Signaling Technology, #80772, 1:1000), anti-MTAP (Cell Signaling Technology, #41585, 1:1000), anti–symmetric dimethylarginine (SYM10; Cell Signaling Technology, #13222, 1:1000), and anti–β-actin (Cell Signaling Technology, #3700, 1:1000). An HRP-conjugated anti-rabbit IgG secondary antibody (Cell Signaling Technology, #7074S) was used. Blot imaging was carried out using the G:BOX Chemi XRQ system with GeneSys software v1.6.5.0 (Syngene). Band intensities were quantified using Gene Tools analysis software.

### Cell Viability Assay

Cell viability was assessed using the CellTiter-Glo® Luminescent Cell Viability Assay (Promega) following the manufacturer’s protocol. 2,500 cells per well were seeded into 96-well plates in 150 μL of culture medium 24 hours prior to drug treatment. Compounds were added using a digital drug printer. On the day of evaluation, plates were equilibrated to room temperature, and 37.5 μL (20% of final volume) of CellTiter-Glo reagent (Promega, G7573) was added to each well. After a 30-minute incubation at room temperature, luminescence was measured using a Spectra Max i3x microplate reader with SoftMax Pro software v7.0.2 (Molecular Devices).

### Synergy analysis

Drug combinations were evaluated in 384-well plates using the same viability assay protocol described above. Growth inhibition (GI) was calculated using the following formula: If T < D0: GI = 100 × (1 − ((Signal − D0) / D0)), If T ≥ D0: GI = 100 × (1 − ((Signal − D0) / (Vehicle − D0))). Here, D0 represents the background luminescence at time 0 (before treatment), and T is the luminescence signal after treatment. Synergistic effects were analyzed using the SynergyFinder Plus web application, applying the Loewe additivity model. ^20^

### Cell Monitoring

Continuous cell monitoring was performed using the Incucyte live-cell imaging system. Cell proliferation was quantified based on % confluence or by using the “adherent cell-by-cell” mode, depending on the experimental context.

### Crystal Violet Colony Formation Assay

Cells were seeded to give a confluency of ∼80% and drugged 6 days. After drugging, the cells were fixed with 10% glutaraldehyde (Fisher Scientific, #BP2547-1) and stained with 0.1% crystal violet (Sigma, V5265-500mL).

### Statistical Analysis

For cell line studies, data analysis was conducted using GraphPad Prism software v10.6.1 (GraphPad Software). For comparisons between experimental groups and controls, Student’s t-tests or Mann–Whitney U tests were used as appropriate. A p-value of <0.05 was considered statistically significant by convention. Fisher’s exact test was used to compare frequency of MTAP loss in specimens from patients with advanced (i.e., stage IV) vs stage I-III NSCLC and with driver-positive vs driver-negative NSCLC. All p-values were based on a two-sided hypothesis and computed using Stata/SE 17.0.

## RESULTS

### Study Population

The tissue dataset (**Figure 1**) was comprised of 243 specimens, including 132 lung resection specimens from patients with stage I-III NSCLC (n=66 stage I; n=39 stage II; n=27 stage III) and 111 specimens from 108 patients with treatment-naïve stage IV NSCLC. The patients included 96 (40%) males and 144 (60%) females (**Supplementary Table 1**). The tissue dataset originated from a broader institutional effort to assess expression of diverse protein biomarkers across a cohort enriched for oncogenic driver alterations.

**Figure 1.**
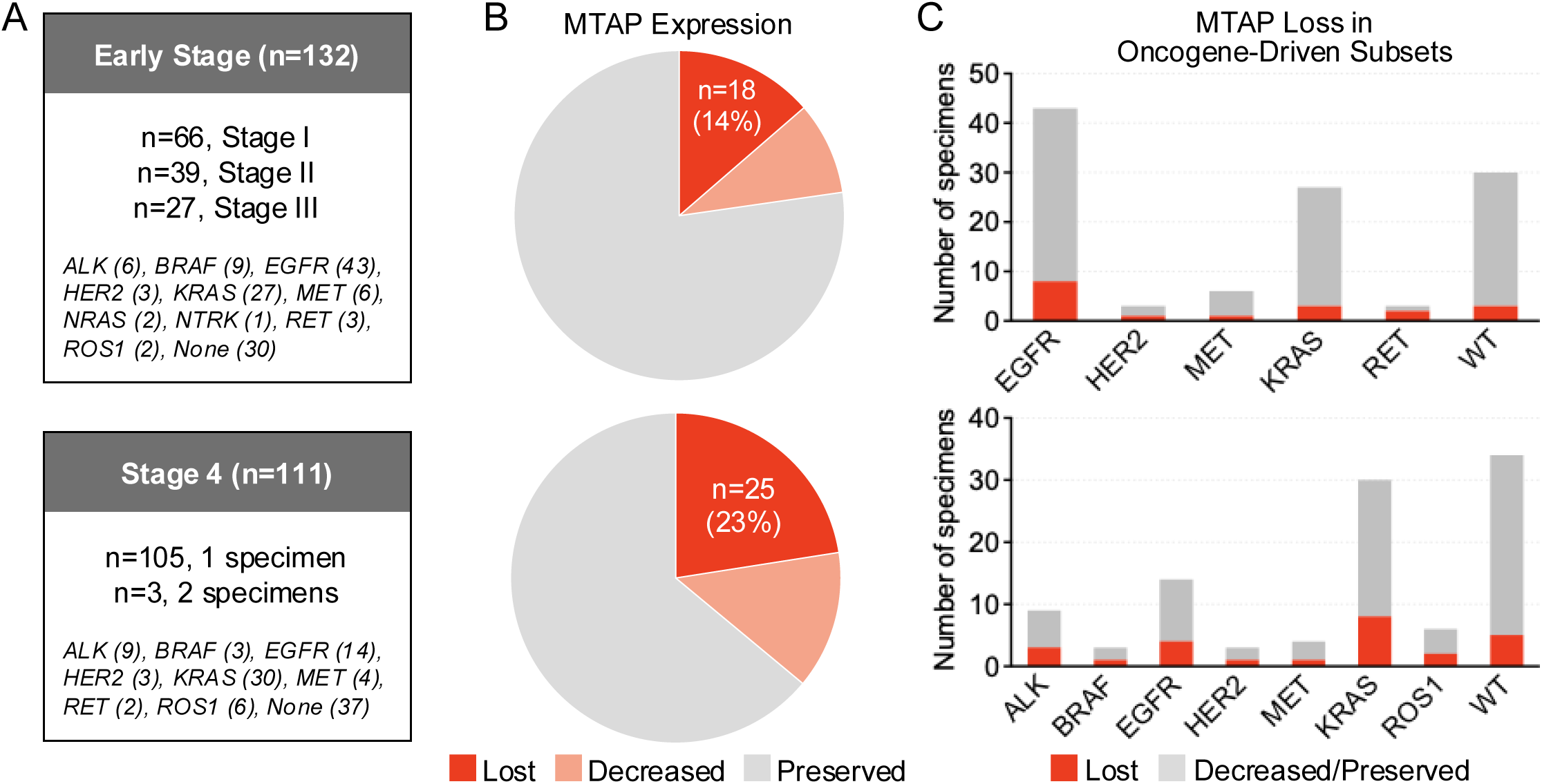
MTAP Expression by Stage and According to Molecular Context. (A) Boxes summarize the specimens included in the early stage and stage 4 datasets, including molecular alterations identified by next-generation sequencing. (B) Pie charts depict MTAP expression in early stage (top) and stage 4 (bottom) as assessed by immunohistochemistry. (C) Bar graphs illustrate prevalence of MTAP loss as assessed by immunohistochemistry in early stage (top) and stage 4 (bottom) specimens delineated by molecular driver status.

#### Early-Stage NSCLC

126 (95%) of 132 specimens were adenocarcinoma histology (**Supplementary Table 1**). The distribution of molecular alterations was as follows: *ALK* fusion (n=6), *BRAF* V600E (n=4), class II-III *BRAF* mutation (n=5), *EGFR* exon 19 deletion (n=13), *EGFR* L858R (n=24, including 1 case concurrent with 768I), atypical *EGFR* mutation only (n=3), *EGFR* exon 20 insertion (n=3), *HER2* alteration (n=2 insertion; n=1 amplification), *KRAS* mutation (n=27), *MET* exon 14 skipping (n=6), *NRAS* mutation (n=2), *NTRK1* fusion (n=1), *RET* fusion (n=3), and *ROS1* fusion (n=2). The remaining 30 specimens did not contain established molecular driver alterations. Paired primary lung tumor and thoracic lymph node were available for 33 of the specimens.

#### Stage IV NSCLC

The stage IV specimens included 111 specimens collected from 108 patients (**Figure 1, Supplementary Table 1)**. Nine specimens were archival lung resection tissue obtained at initial diagnosis of early-stage lung cancer. All specimens were collected prior to initiation of systemic therapy. Adenocarcinoma histology was identified in 89 (82%) specimens. The following molecular driver alterations were detected in the 108 specimens (excluding repeat biopsies from 3 patients): *ALK* fusion (n=9), *BRAF* V600E (n=1), class II-III *BRAF* mutation (n=2), *EGFR* exon 19 deletion (n=5), *EGFR* L858R (n=6), *EGFR* exon 20 insertion (n=3), *HER2* mutation (n=3), *KRAS* mutation (n=30), *MET* alteration (n=3 *MET* skipping, n=1 *MET* amplification), *RET* fusion (n=2), and *ROS1* fusion (n=6). Specimens from 37 patients did not harbor identifiable established molecular driver alterations.

### Prevalence of MTAP loss in NSCLC Specimens

By IHC, completely preserved MTAP expression was identified in 173 (71%) of the 243 specimens, including 102 (77%) early-stage NSCLCs and 71 (64%) specimens from patients with metastatic NSCLC (**Figure 1)**. Decreased or absent MTAP expression was observed in 70 (29%) of the 243 specimens. Forty-three (18%) specimens had complete loss of MTAP expression, including 18 (14%) early-stage NSCLCs (of which one also had *CDKN2A* loss) and 25 (23%) specimens from patients with metastatic NSCLC (**Figure 1B-1C**). The 25 specimens with MTAP loss from the metastatic cohort originated from 22 patients such that three patients had consistent MTAP loss across two distinct specimens. Twenty-seven (11%) specimens had heterogeneous/dichotomous MTAP loss or decreased expression without complete loss (**Figure 1**). Representative images of preserved, heterogeneous, and absent MTAP expression are presented in **Supplementary Figure 1.**

To ascertain whether MTAP loss was enriched in NSCLCs harboring driver alterations, we compared the prevalence of MTAP loss in driver-negative vs driver-positive specimens in the early-stage and metastatic groups (**Figure 2)**. For this analysis, only complete absence of MTAP expression was considered MTAP loss. For the three patients with multiple biopsies with MTAP loss, only one biopsy was included. Among 18 early-stage NSCLCs with complete loss, 15 (83%) contained driver alterations, i.e., three (10%) of 30 driver-negative versus 15 (15%) of 102 driver-positive specimens exhibited MTAP loss (p=0.763, **Figure 2A-2B**). In the stage IV dataset, complete MTAP loss was observed in 5 (14%) of 37 driver-negative tumors versus 17 (24%) of 71 driver-positive tumors (p=0.314, **Figure 2C-D**). Within the *EGFR* and *KRAS* tumors, frequency of MTAP loss was compared for early-stage versus metastatic NSCLCs. In *EGFR*-mutant specimens, MTAP loss occurred in eight (19%) of 43 early-stage vs four (29%) of 14 metastatic specimens (p=0.463). MTAP loss was observed in three (11%) of 27 early-stage vs six (20%) of 30 metastatic *KRAS-*mutant NSCLCs (p=0.476). Prevalence of MTAP loss with specific driver alterations is presented in **Supplementary Table 2 and Figure 1C**. Overall, our findings suggest that a subset of NSCLCs across disease stages and molecular contexts harbors MTAP loss, with highest prevalence in advanced tumors.

**Figure 2.**
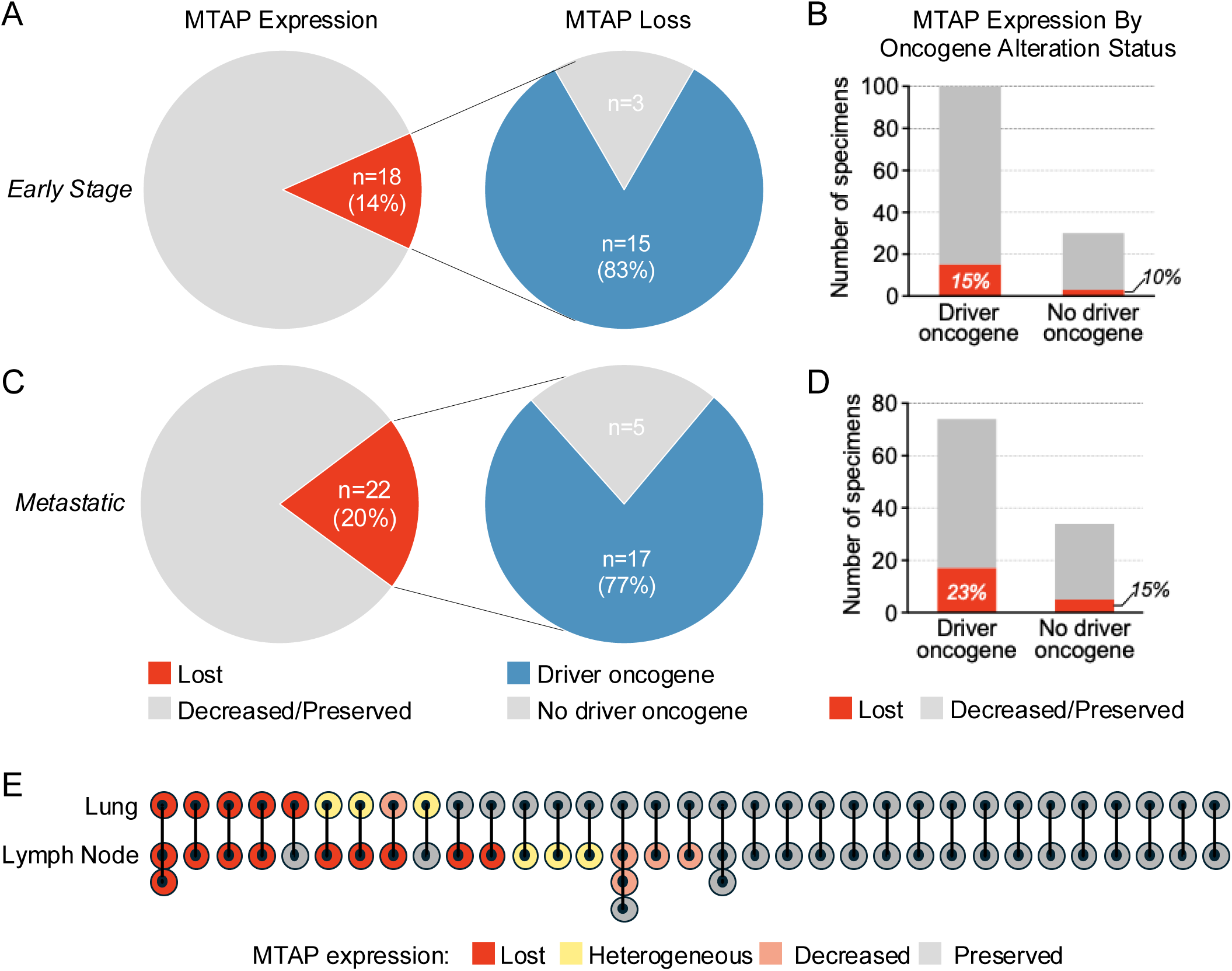
MTAP loss according across disease stage, according to driver alteration status, and in primary tumor versus involved thoracic lymph nodes. (A) Pie charts depict MTAP expression as assessed by immunohistochemistry in entire early stage cohort and in tumors with and without driver alterations. (B) Bar graphs summarize proportion of early stage lung cancer specimens with MTAP loss as assessed by immunohistochemistry among tumors with and without driver oncogenes. (C) Pie charts depict MTAP expression as assessed by immunohistochemistry in entire metastatic lung cancer cohort and in tumors with and without driver alterations. (D) Bar graphs summarize proportion of metastatic lung cancer specimens with MTAP loss as assessed by immunohistochemistry among tumors with and without driver oncogenes. (E) Figure illustrates MTAP expression as assessed by immunohistochemistry in lung primary compared to associated involved thoracic lymph nodes. Heterogeneous indicates mixed population with some cells exhibiting loss whereas other adjacent cells demonstrate preserved MTAP expression.

Next, we evaluated heterogeneity of MTAP loss by examining MTAP expression in paired lung resection tissue and involved lymph nodes from 33 patients (**Figure 2E**). We also explored the utility of small biopsies for determining MTAP status by comparing pre-surgical lung biopsy findings to resection specimens from 29 patients. Presurgical biopsy reliably captured MTAP status in the resection specimen in 28 (97%) of 29 cases. Concordance between MTAP status in resected lung and nodal tissue was seen in 20 (61%) cases (**Figure 2E**). Discordance between MTAP expression in resected lung and nodal tissue was observed in 13 instances, including one instance in which MTAP loss was confined to the lung tissue and 8 instances in which MTAP expression was lost or reduced in node but completely preserved in lung tissue. In the remaining 4 cases, the degree of discordance was less pronounced. Thus, reliance on lung biopsies may underestimate the prevalence of MTAP loss.

### MTAP-Deficient Oncogene-Driven NSCLC Cell Lines Have Variable Sensitivity to PRMT5 Inhibition

To investigate the sensitivity of MTAP-deficient oncogene-driven NSCLCs to MTA-cooperative PRMT5 inhibitors, we first screened a panel of 63 oncogene-driven NSCLC cell lines (17 *EGFR*-mutant, 22 *KRAS*-mutant, 21 *ALK*-rearranged, 2 *ROS1*-rearranged, and 1 *RET*-rearranged, **Figure 3A, Supplementary Figure 2A, Supplementary Table 3**) to identify those with MTAP protein loss. To enrich for MTAP loss, we restricted the screen to cell lines with known *CDKN2A* alterations or those for which CDKN2A testing had not been previously performed. Genomic annotations for publicly available cell lines were obtained from the Cancer Cell Line Encyclopedia (CCLE). For cell lines established at MGH, genomic status of the original patient tumor was determined by NGS-based copy number profiling. For comparison, we also examined nine cell lines without *CDKN2A* alterations. Loss of CDKN2A (p16^INK4A^) protein expression was confirmed in 53 models, with concurrent complete loss of MTAP protein expression in 22 models (42%): 9 *EGFR*-mutant, 7 *KRAS*-mutant, and 6 *ALK*-rearranged (**Figure 3A, Supplementary Figure 2A-C**).

**Figure 3.**
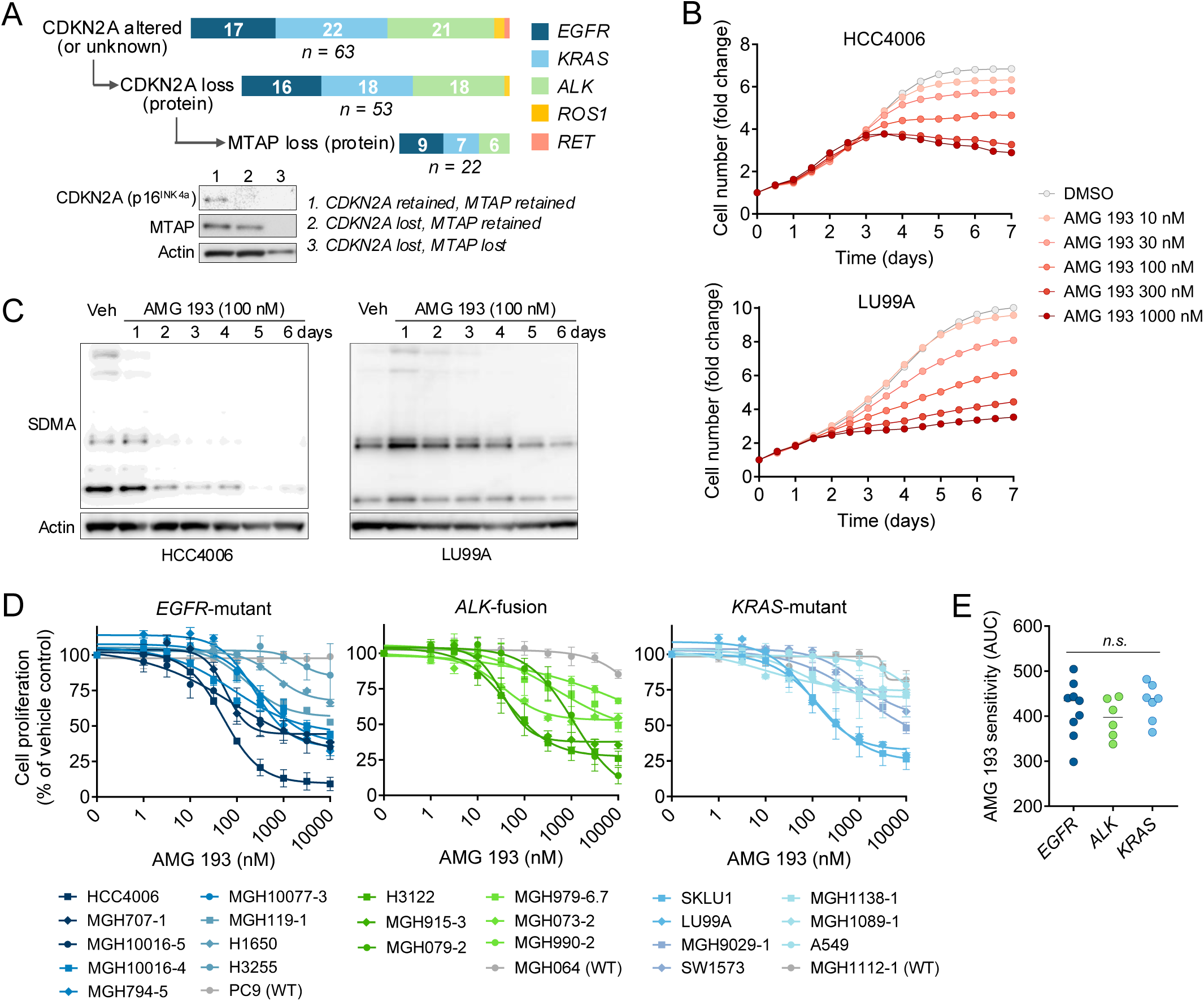
Sensitivity of MTAP-deficient, oncogene-driven NSCLC cell lines to PRMT5 inhibition. (A) Oncogene-driven NSCLC cell lines were screened for CDKN2A and MTAP protein expression. Lower panel shows representative CDKN2A and MTAP status as determined by western blotting. (B) Time course of cell proliferation following treatment of MTAP-deficient cell lines with increasing concentrations of AMG 193 as determined by live-cell imaging. (C) Western-blotting of symmetric dimethylarginine (SDMA) levels following AMG 193 treatment in MTAP-deficient models. (D) Cell proliferation of 22 MTAP-deficient NSCLC cell lines was assessed by CellTiter-Glo following AMG 193 treatment for 10 days. (E) Comparison of AUC values from AMG 193 cell proliferation assays (Panel D), stratified by oncogenic driver mutations.

To evaluate sensitivity to PRMT5 inhibition, MTAP-deficient cell lines were treated with increasing concentrations of AMG 193, an MTA-cooperative PRMT5 inhibitor currently being evaluated in clinical trials.^14,15^ We observed suppression of proliferation beginning 2-4 days after addition of drug (**Figure 3B, Supplementary Figure 3A**), with the effect becoming more apparent over time (**Supplementary Figure 3B**), consistent with prior studies of PRMT5 inhibitors.^21,22,15^ Suppression of proliferation coincided with a progressive time-dependent decrease in symmetric dimethylarginine (SDMA) levels, a direct readout of PRMT5 enzymatic function (**Figure 3C**). Comparing the effects of AMG 193 across the cohort of 22 MTAP-deficient *EGFR*-mutant, *KRAS-*mutant, and *ALK*-positive cell lines, we observed variable activity with some MTAP-deficient cell lines showing > 90% suppression of proliferation while others showed limited sensitivity (similar to MTAP wild-type cell lines) (**Figure 3D**). Similar relative drug sensitivity profiles were observed with other clinical-grade MTA-cooperative PRMT5 inhibitors—TNG462 and MRTX1719—albeit with differences in absolute potency (**Supplementary Figure 3C-E**). No association between driver oncogene and drug response was observed in our cell line cohort (**Figure 3E**). These results demonstrate that a subset of oncogene-addicted NSCLCs with concurrent MTAP loss are sensitive to PRMT5 inhibition, largely independent of the specific oncogenic driver.

### SDMA as a Pharmacodynamic, but not Predictive, Marker of PRMT5 Inhibition

We next examined whether baseline or dynamic changes in SDMA levels correlate with sensitivity to PRMT5 inhibition. Baseline SDMA levels were lower in MTAP-deficient compared to MTAP wild-type cell lines (**Supplementary Figure 4A**), consistent with prior reports.^23,24^ However, when stratified by AMG 193 sensitivity (defined as an IC₅₀ < 1 μM), baseline SDMA levels did not distinguish sensitive from insensitive MTAP-deficient cell lines (**Figure 4A**). Next, we examined treatment-induced changes in SDMA. AMG 193 treatment reduced SDMA in both MTAP-deficient and WT cell lines; however, significantly higher concentrations were required for equivalent SDMA suppression in MTAP-WT cell lines (**Figure 4B-C, Supplementary Figure 4B**). The magnitude or potency of SDMA reduction did not correlate with cellular sensitivity to AMG 193 (**Figure 4D, Supplementary Figure 4C**). These results indicate that SDMA levels are a reliable dynamic pharmacodynamic marker of PRMT5 inhibition but do not predict therapeutic efficacy amongst oncogene-driven MTAP-deficient NSCLCs.

**Figure 4.**
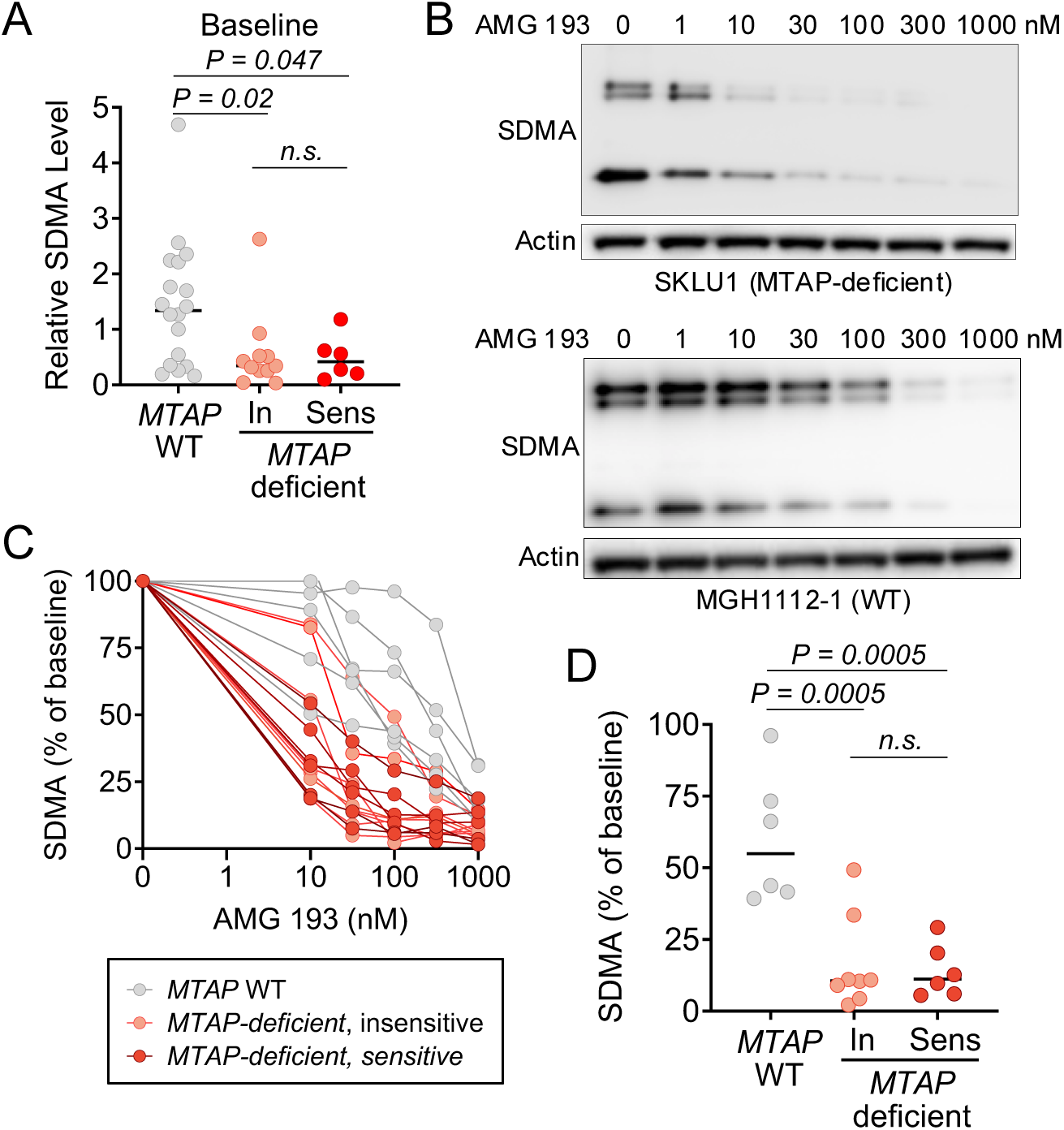
SDMA is a pharmacodynamic but not predictive biomarker of PRMT5 inhibition in NSCLC. (A) Baseline SDMA levels in MTAP-deficient and MTAP wild-type oncogene-driven NSCLC cell lines, stratified by sensitivity to AMG 193, which defined as an IC₅₀ < 1 μM. (B) Dose-dependent reduction in SDMA levels following AMG 193 treatment in representative MTAP-deficient (SKLU1) and MTAP wild-type (MGH1112-1) cell lines. Cells were treated with indicated concentrations of AMG 193 for six days prior to harvesting for western blot analysis. (C) Quantification of SDMA suppression across cell lines following AMG 193 treatment, illustrating differences in potency between MTAP-deficient and MTAP wild type backgrounds. (D) Comparison of SDMA suppression after treatment with 100 nM AMG 193 (six days) between sensitive and insensitive MTAP-deficient cell lines, and MTAP wild-type cell lines.

### PRMT5 Inhibition Enhances the Efficacy of Oncogene-Targeted Therapies

As PRMT5 regulates mechanisms orthogonal to kinase signaling cascades activated by EGFR, KRAS, or ALK, we evaluated whether PRMT5 inhibition enhances the efficacy of oncogene-directed targeted therapies. We treated MTAP-deficient cell lines with AMG 193 in combination with genotype-matched targeted therapies (EGFR: osimertinib, KRAS: RMC-6236, ALK: lorlatinib). Across oncogenic drivers, dual drug treatment resulted in consistently greater suppression of proliferation than either agent alone (**Figure 5A, Supplementary Figure 5A**). When comparing the kinetics of drug response, oncogene-targeted therapies exerted immediate effects, whereas the effects of AMG 193 alone or in combination were more pronounced after several days, consistent with the delayed pharmacodynamic effects of PRMT5 inhibition (**Figure 5B**). Next, we performed 2x2 dose-response matrices to determine whether the combination effect was synergistic or additive. While synergy analysis revealed modest variability across cell lines and dosing conditions with some combinations exhibiting weak synergy, most interactions were additive (**Figure 5C, Supplementary Figure 5B-E**). These results suggest that the benefit of PRMT5 inhibition in combination with targeted therapies is largely due to independent action of each drug.

**Figure 5.**
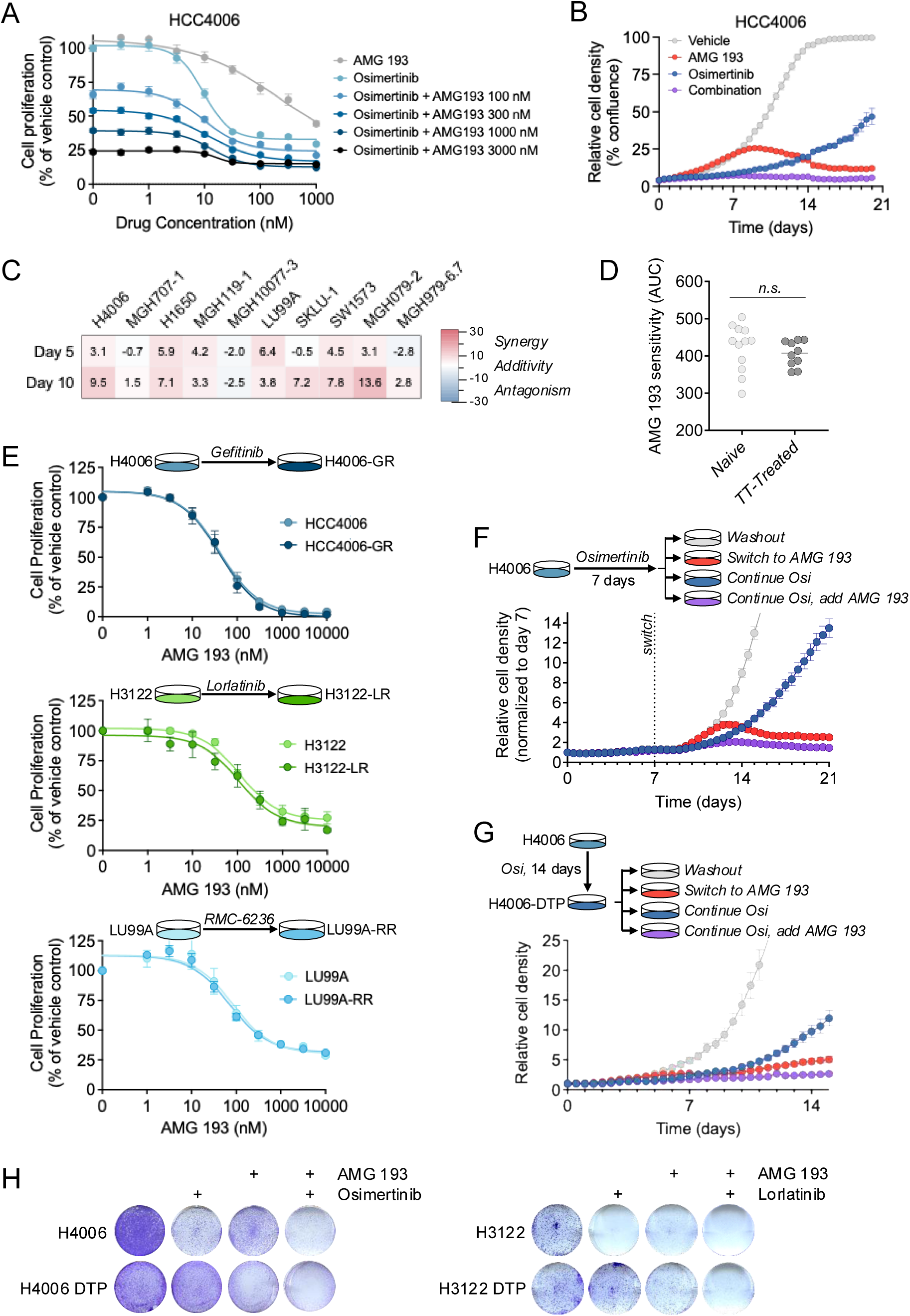
PRMT5 inhibition augments targeted therapy efficacy. (A) Proliferation assays of MTAP-deficient HCC4006 cells treated with AMG 193 for six days in combination with 1 μM osimertinib. (B) Time-course of HCC4006 cell proliferation assessed by live-cell imaging following treatment with AMG 193, osimertinib or their combination. (C) Loewe synergy scores of MTAP-deficient, oncogene-driven NSCLC cell lines were calculated from cell proliferation assays performed cells treated with the combination of TKI (osimertinib – EGFR, lorlatinib – ALK, RMC-6236 – KRAS) and AMG 193 for 5 or 10 days. (D) AMG 193 sensitivity (AUC of cell proliferation dose-response assays) in cell lines derived from targeted therapy–naïve versus previously treated tumors. (E) Cell proliferation of matched parental and resistant cell line pairs (HCC4006/HCC4006-GR, H3122/H3122-LR, and LU99A/LU99A-RR) following treatment with AMG 193 for six days. (F) Time course of HCC4006 cell proliferation assessed by live-cell imaging upon treatment with 1 μM osimertinib followed by switch to or addition of 100 nM AMG 193. (G) HCC4006 cells were treated with osimertinib for 14 days to establish DTP cells, the treated with the indicated drugs. Cell proliferation was assessed by live-cell imaging. (H) Colony formation assays demonstrating enhanced efficacy of AMG 193 and TKI combination in both parental and DTP conditions. DTP populations were generated by treatment with osimertinib (HCC4006) or lorlatinib (H3122) for 14 days prior to the experiment.

### Sensitivity to PRMT5 Inhibition is Independent of Prior TKI Exposure

To determine whether dependency on PRMT5 evolves over the clinical course of treatment, we examined whether a history of prior targeted therapy exposure affected AMG 193 sensitivity. In our cohort of *EGFR*-mutant and *ALK*-positive models, five of nine and five of six cell lines, respectively, were generated from patients whose cancer had progressed on EGFR or ALK targeted therapies (**Supplementary Table 3**). None of the seven *KRAS*-mutant models had been exposed to prior targeted therapy. Comparison of targeted therapy-naïve (n = 12) and resistant (n = 10) models revealed no significant differences in AMG 193 sensitivity (**Figure 5D**), suggesting that development of acquired resistance to targeted therapies does not impact PRMT5 dependence. To confirm these findings, we compared the sensitivity of matched targeted therapy-naïve and resistant cell line pairs: HCC4006/HCC4006-GR, H3122/H3122-LR, and LU99A/LU99A-RR (**Supplementary Figure 6A**).^18,19^ Resistant cell lines showed similar AMG 193 sensitivity as the parental cell line from which they were derived (**Figure 5E, Supplementary Figure 6B**). Collectively, these data indicate that AMG 193 activity is not impacted by prior targeted therapy treatment and suggest that PRMT5 inhibitor sensitivity is independent of other mechanisms of acquired resistance to oncogene-targeted therapies.

### PRMT5 Inhibition Suppresses Outgrowth of TKI-Persistent Cells

The observations that PRMT5 inhibitors act independently of oncogene-targeted therapies and that PRMT5 dependency is sustained in the context of TKI resistance motivated us to explore whether PRMT5 dependency might be a vulnerability of MTAP-deficient drug tolerant persister (DTP) cells, which we and others have shown can serve as the precursors for outgrowth of fully resistant tumor clones.^25,26^ In parental HCC4006 cells, osimertinib initially suppressed cell proliferation, followed by regrowth (**Figure 5F**). Adding AMG 193 to osimertinib on day 7 suppressed the outgrowth of drug-tolerant persister cells. We corroborated these findings by first establishing HCC4006 DTP populations (by treating with 1 μM osimertinib for 2 weeks) and then treating with AMG 193 alone or in combination with osimertinib. AMG 193 alone suppressed persister outgrowth relative to continued osimertinib, with further suppression when combined with continued osimertinib (**Figure 5G**). Similar results were observed in colony formation assays, which confirmed the enhanced effect of the combination in both parental and persister settings (**Figure 5H**). These results suggest that targeting dependencies such as PRMT5 that are retained in resistant and drug tolerant persister states may represent a strategy for pre-empting the development of drug resistance.

## DISCUSSION

MTAP loss is an emerging therapeutic target for which there are promising MTA-selective PRMT5 inhibitors under development. Reliably identifying MTAP loss is critical for recruitment to ongoing trials, as are studies that can help inform sequencing strategies for patients with NSCLCs harboring both MTAP loss and actionable oncogenic driver alterations. In this manuscript, we assessed MTAP expression in 243 NSCLC specimens and evaluated the activity of PRMT5 inhibitors in patient-derived cell lines reflecting three prevalent molecular contexts. Through these studies, we demonstrate that MTAP loss occurs in 18% of non-squamous NSCLC, including one-fourth of metastatic oncogene-driven NSCLCs, with a subset of such tumors demonstrating sensitivity to PRMT5 inhibitor monotherapy that was independent of prior TKI exposure and enhanced by the addition of TKI therapy. Our findings reinforce the necessity of diagnostic approaches that interrogate both activated oncogenes and therapeutically relevant copy losses to expand treatment opportunities for NSCLC patients.

We assessed MTAP expression in adenocarcinomas across disease stages and anatomic sites. MTAP loss was identified in 18% of the 243 specimens (17% of 240 patients), including 14% of early-stage and 20% of metastatic NSCLCs. Although the higher rate of MTAP loss in advanced disease was not statistically significant, the relative increase in prevalence of MTAP loss suggests against relying on MTAP status in resection specimens in cases where metastatic relapse has occurred. This notion is further supported by the 39% discordance observed between MTAP expression in nodal metastases versus primary lung tumors, with 24% of cases exhibiting decreased MTAP expression exclusively in lymph nodes. Our dataset similarly suggests against using CDKN2A status derived by NGS to deduce MTAP status, as *CDKN2A* alterations were rarely reported in our MTAP loss cohort. Our *CDKN2A* analysis was confined to early-stage tumors evaluated with Genexus OPA due to lack of coverage for *CDKN2A* copy changes in the assays used for metastatic specimens. It is likely that the lower detection rate for *CDKN2A* alterations in our analysis reflects Genexus OPA performance limitations as recently reported by others,^3^ as an independent analysis of frequency of *CDKN2A* alterations in 470 consecutive NSCLCs analyzed using Genexus OPA at our institution demonstrated *CDKN2A* loss in only 0.6% (n=1/166) of early-stage and 6% (n=18/304) of metastatic lung adenocarcinomas, respectively (**Supplementary Figure 7)**. This lower-than-expected rate of *CDKN2A* loss highlights the challenges of NGS versus IHC for querying copy number losses. Indeed, identifying copy number variation using bulk NGS relies on specimen quality, tumor purity, and assay specifications including breadth of genomic coverage.

We selectively evaluated PRMT5 inhibition in the three most prevalent actionable subsets of NSCLC: *EGFR* mutations, *KRAS* mutations, and *ALK* fusions. Consistent with clinical experience for PRMT5 inhibitors,^15,21,22^ PRMT5 inhibitor monotherapy demonstrated robust antiproliferative activity in a subset of oncogene-positive, MTAP-deficient cell lines. We found that monotherapy activity of PRMT5 inhibitors was independent of the driver alteration and did not correlate with baseline or dynamic SDMA levels. Thus, given the imperfect nature of MTAP as a predictive biomarker, there is unmet need for more incisive biomarkers that better delineate vulnerability to PRMT5 inhibition. The imperative to identify biomarkers for PRMT5 inhibition is rationalized by response kinetics of this class of drugs, specifically the delayed onset of responses such that patients incur drug exposures and toxicities while awaiting effect.

Therapeutic strategies for *EGFR-*mutant and *ALK*-positive NSCLC continue to evolve with recent adoption of combination strategies for the former and prioritization of highly efficacious TKIs for the latter. These treatment modifications have created a post-progression space characterized by a lack of viable personalized therapies for TKI-resistant tumors. In contrast, the modest clinical activity of approved G12C inhibitors in *KRAS*-mutant NSCLC has confined targeted therapy to the later-line setting and generated interest in developing combinations that augment efficacy. In our study, AMG 193 was active in both sensitive and resistant models, supporting utility of this therapy across treatment contexts and in tumors with diverse signaling dependencies. The context-agnostic efficacy of PRMT5 inhibitor monotherapy raises questions regarding the optimal time for introducing PRMT5 inhibition in patients with oncogene-driven MTAP-deleted NSCLC, specifically whether to pursue upfront use or reserve treatment for TKI-resistant tumors. In cell lines, we observed additive antiproliferative activity with the combination of a PRMT5 inhibitor and TKI that surpassed the activity of monotherapy, potentially justifying incorporating PRMT5 inhibitors into the initial treatment course to complement TKI therapy. The argument for earlier use is bolstered by our finding that PRMT5 inhibition restricted the outgrowth of persister cells and retrospective data from other groups suggesting decreased durability of first-line targeted therapies in patients with tumors with MTAP loss.^5^

Our study has several limitations. First, our cohort was predominantly comprised of adenocarcinoma to intentionally enrich for driver oncogenes. Second, the assessment of *CDKN2A* co-alterations was limited by NGS assay performance.^3^ Third, while the overall cohort size of 243 specimens was large, the number of specimens for each molecular subset was small and, thus, restricted our ability to confidently determine the prevalence of *MTAP* loss within distinct subsets. Fourth, we specifically limited our IHC analysis to clinical specimens that had not been exposed to therapeutic selective pressure from TKI therapy; therefore, our manuscript is not positioned to discuss acquired MTAP loss. Finally, the advanced stage and early-stage NSCLC datasets were fully independent precluding us from assessing for acquisition of MTAP loss at metastatic relapse.

In summary, we identified MTAP loss in 20% of oncogene-driven metastatic NSCLC and observed antiproliferative activity of monotherapy with MTA-cooperative PRMT5 inhibitors in a subset of driver-positive MTAP-deleted NSCLC, including TKI-resistant tumors. The enhanced efficacy and persister cell suppression imparted by the combination of TKI + PRMT5 inhibitor relative to either therapy alone provides rationale for exploring combinations as initial therapy.

## Acknowledgements

We would like to thank the patients who provided their specimens for research to support this work and Andrew Do for his assistance with data collation.

---

**Supplementary Figure 1.**
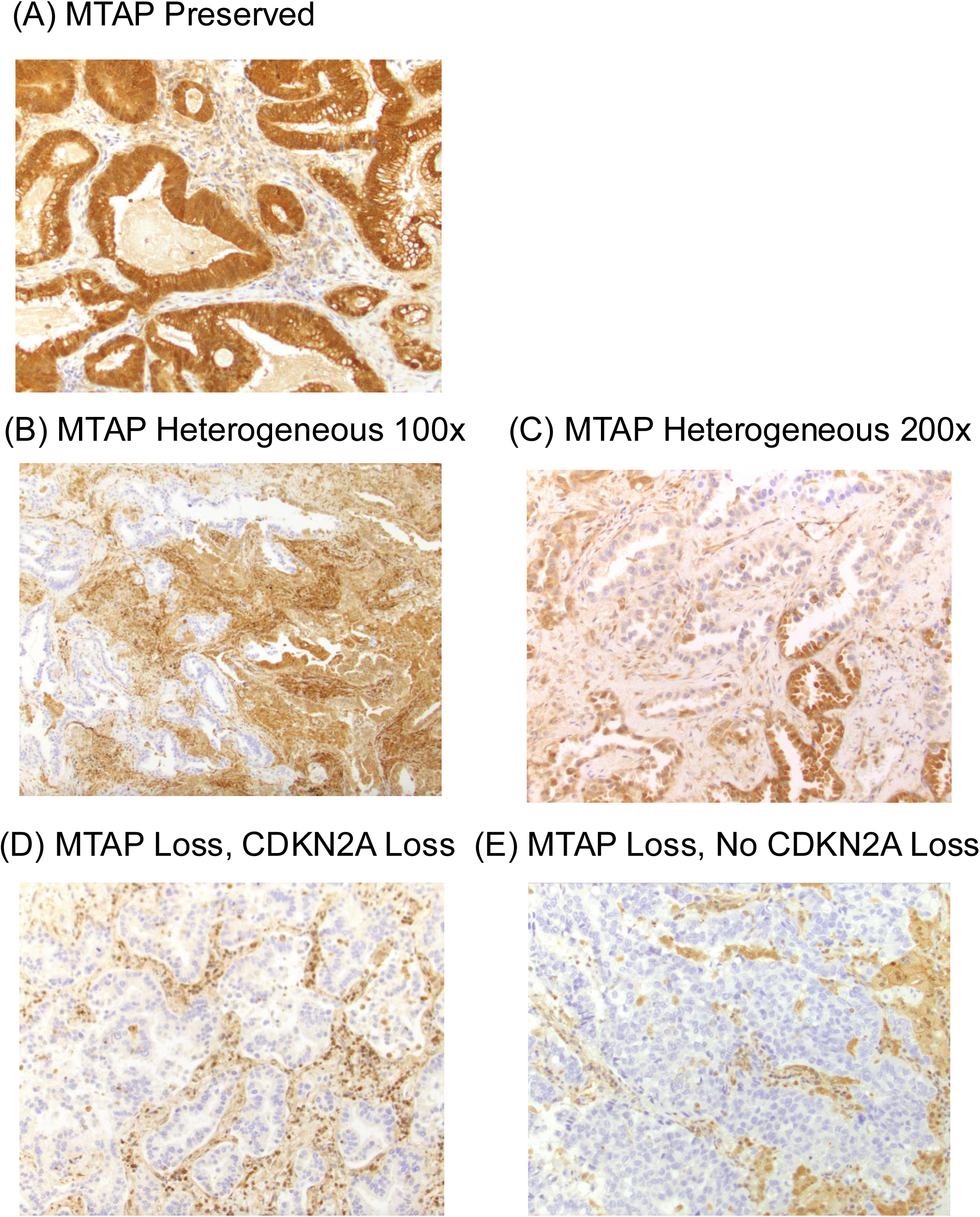
MTAP Expression in Tissue Specimens. Images depict representative specimens with (A) preserved MTAP expression (B) heterogeneous MTAP expression at 100x magnification (C) heterogeneous MTAP expression at 200x magnification (D) loss of MTAP expression with concurrent CDKN2A loss (E) loss of MTAP expression without concurrent CDKN2A loss, as assessed by immunohistochemistry.

**Supplementary Figure 2.**
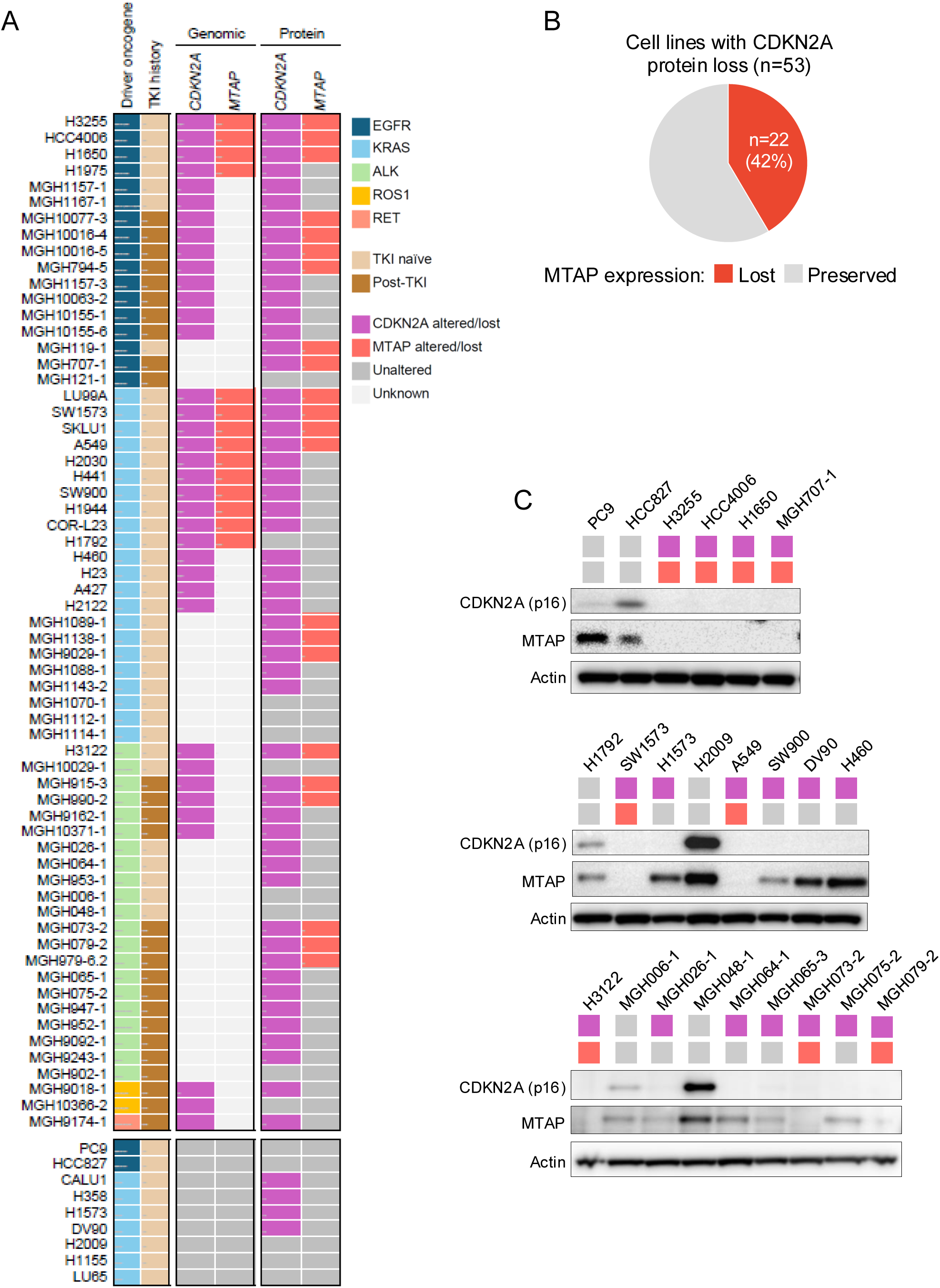
Genomic and protein-level characterization of NSCLC cell lines. (A) Genomic annotation of 63 oncogene-driven NSCLC cell lines, grouped by driver alterations (EGFR, KRAS, ALK, ROS1, RET). Driver mutation status was obtained from CCLE or local genomic profiling. (B) Western blotting of CDKN2A p16^INK4A^ and MTAP protein expression across selected NSCLC cell lines.

**Supplementary Figure 3.**
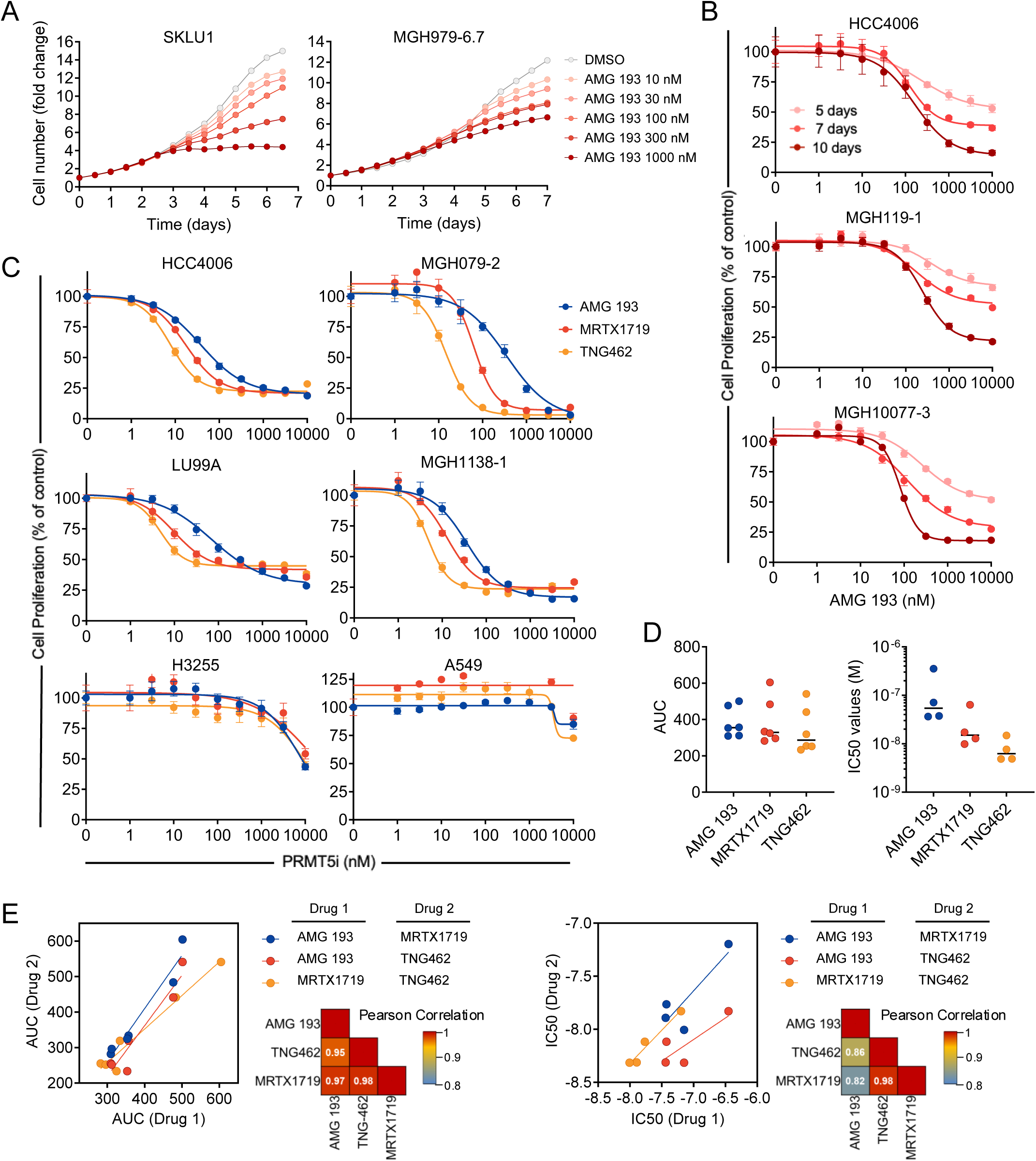
Time- and dose-dependent responses to PRMT5 inhibitors in MTAP-deficient NSCLC cell lines. (A) Time course analysis of cell proliferation following AMG 193 treatment in representative MTAP-deficient models. Inhibition became evident at 2–4 days and increased over time. (B) Prolonged exposure to AMG 193 results in enhanced suppression of proliferation (CellTiter-Glo) in MTAP-deficient cell lines, consistent with delayed pharmacodynamic effects. (C–E) Comparison of inhibition of cell proliferation (AMG 193) in MTAP-deficient NSCLC cell lines treated with the PRMT5 inhibitors AMG193, TNG462, and MRTX1719 for six days.

**Supplementary Figure 4.**
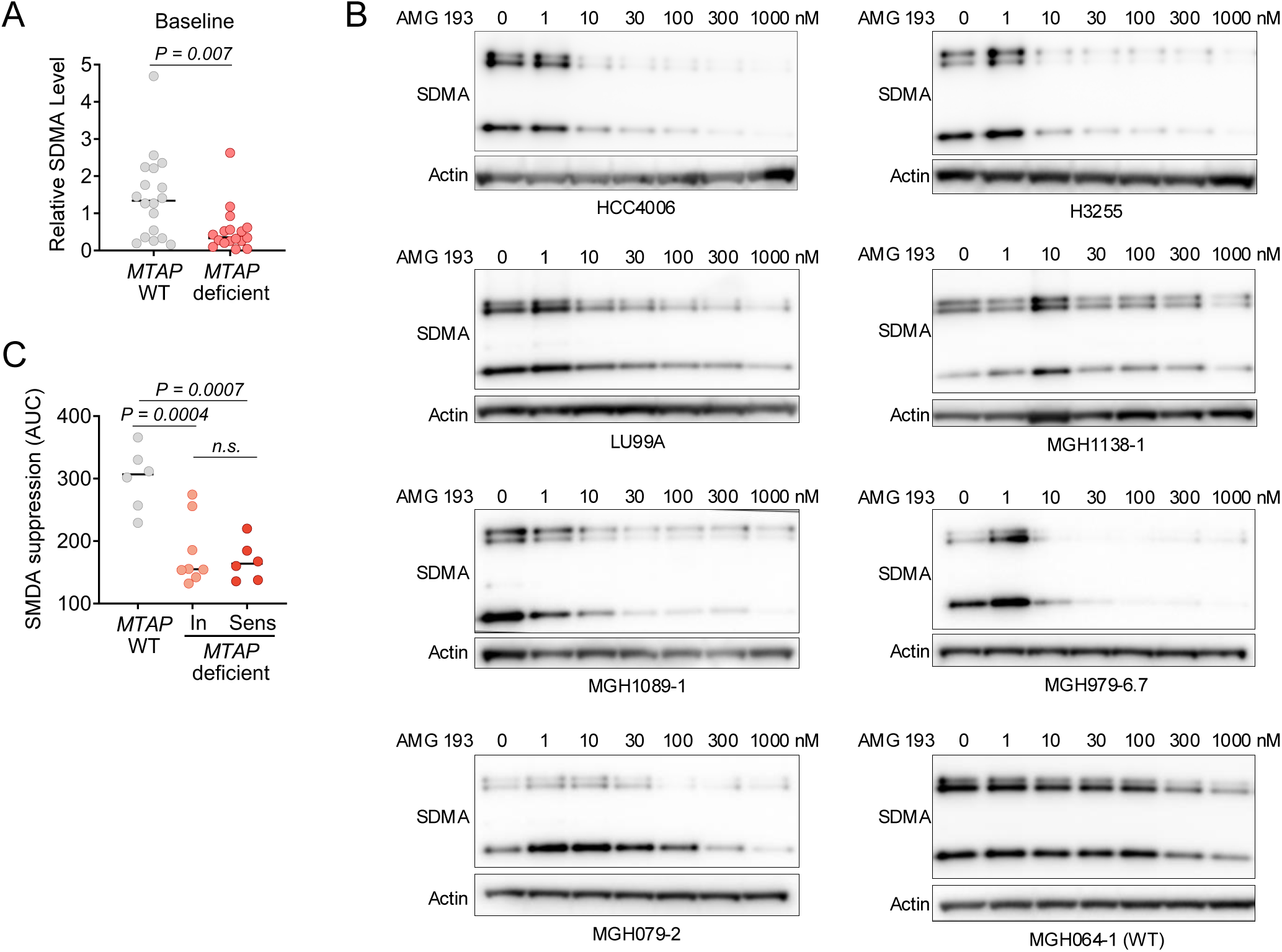
SDMA levels as a pharmacodynamic marker of PRMT5 inhibition. (A) Baseline SDMA levels in MTAP-deficient and wild-type NSCLC cell lines as determined by western blotting. (B) Dose-dependent suppression of SDMA in MTAP-WT and MTAP-deficient cell lines after treatment with AMG 193 for six days. (C) SDMA suppression (AUC of dose response shown in Figure 4C), stratified by insensitive versus sensitive MTAP-deficient cell lines, MTAP wile-type cell lines.

**Supplementary Figure 5.**
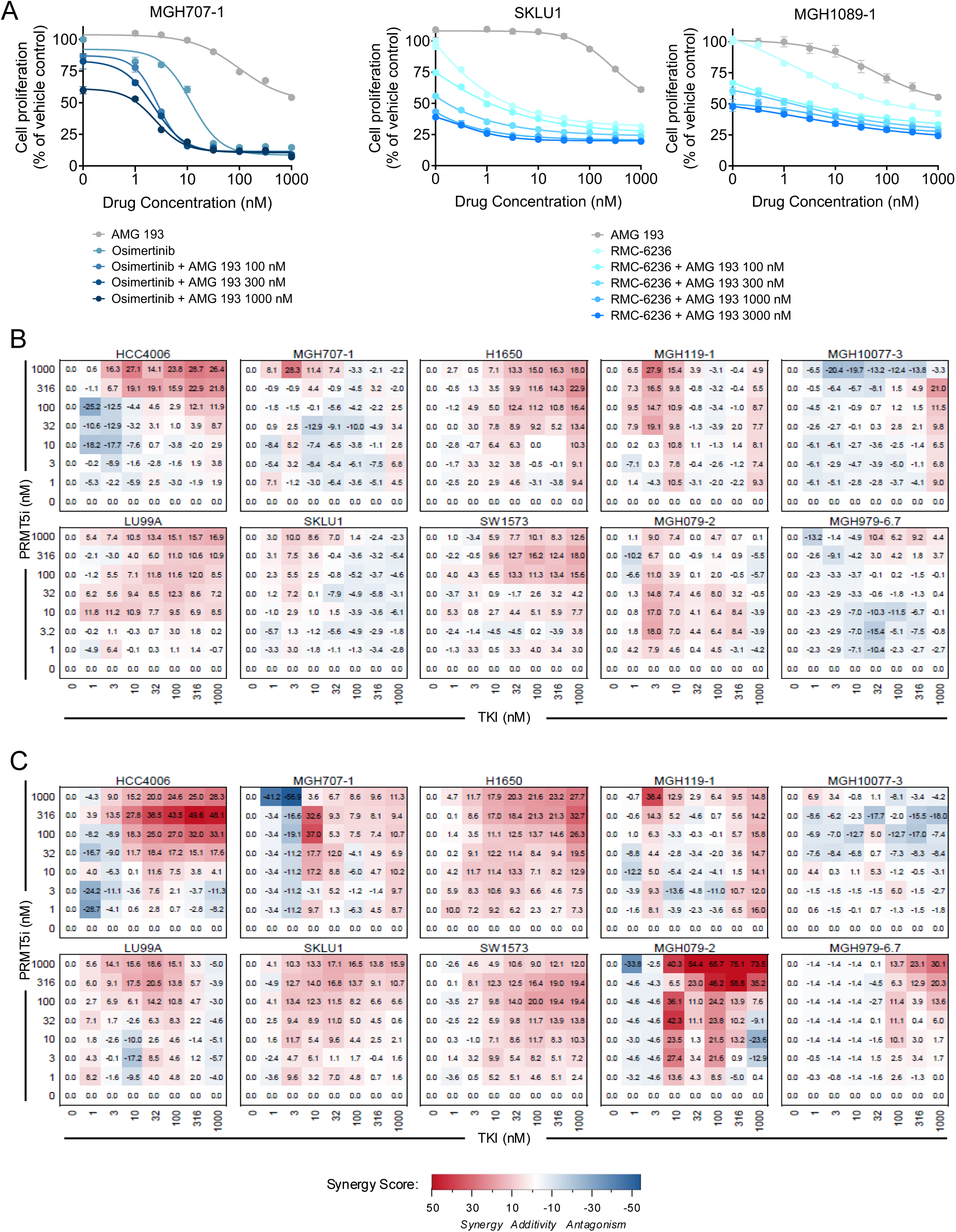
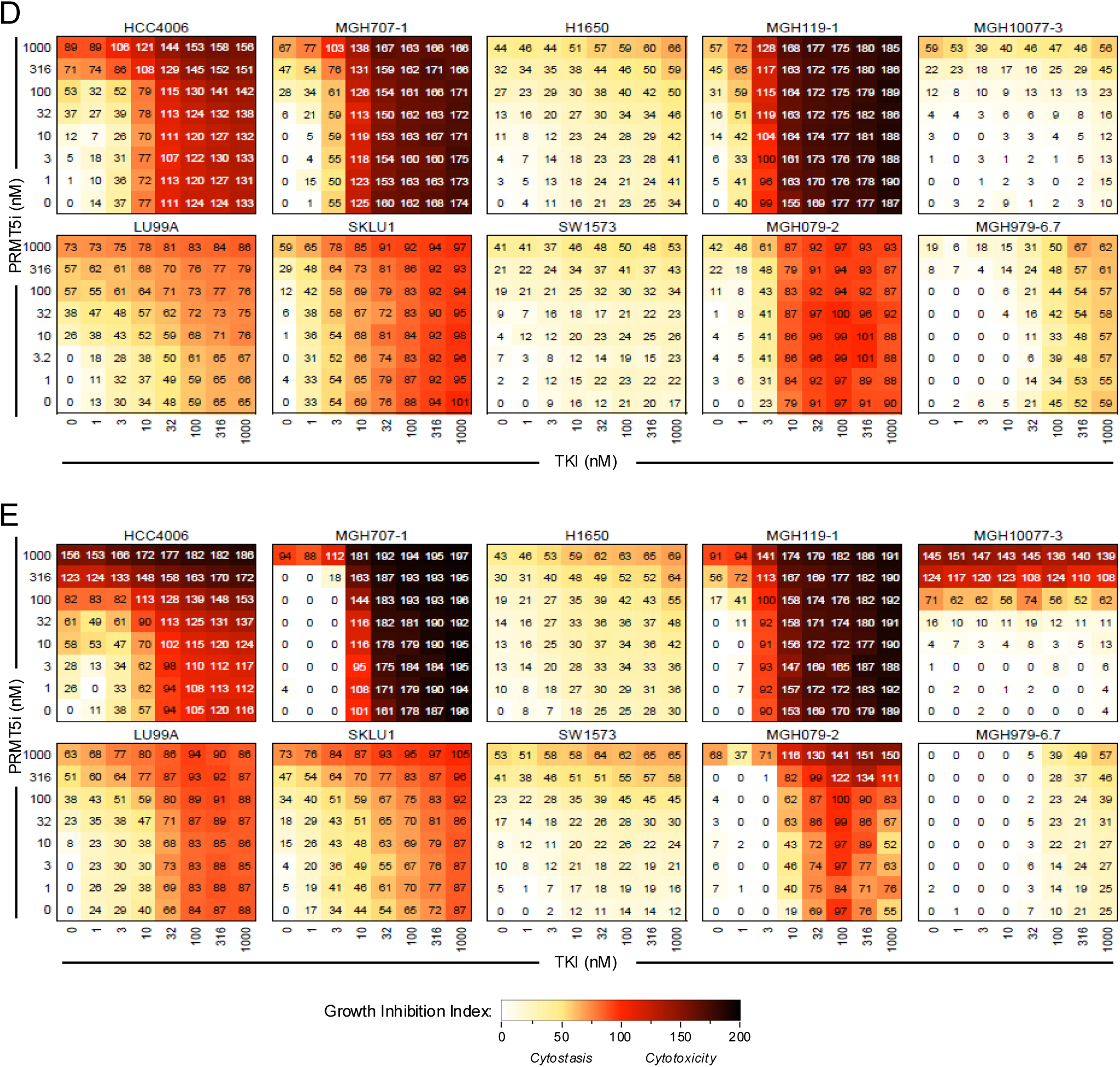
Combination effects of PRMT5 inhibition and targeted therapy. (A) Proliferation assays of MTAP-deficient MGH707-1, SKLU1 and MGH1089-1 cells treated with AMG 193 for six days in combination with 1 μM osimertinib or 300 nM RMC-6236. (B–C) 2×2 dose–response matrices and Loewe synergy scores for AMG 193 combined with targeted therapies after 5 (B) or 10 (C) days of treatment. Most combinations showed additive effects, with some variability in synergy at individual dose levels observed across models. (D–E) Growth inhibition index values for AMG 193 combined with targeted therapies after 5 (D) or 10 (E) days of treatment. 0 = no proliferation effect (value of untreated cells at end of experiment), 100 = cytostasis (no change from day 0), 200 = cytotoxicity (no viability signal at end of experiment).

**Supplementary Figure 6.**
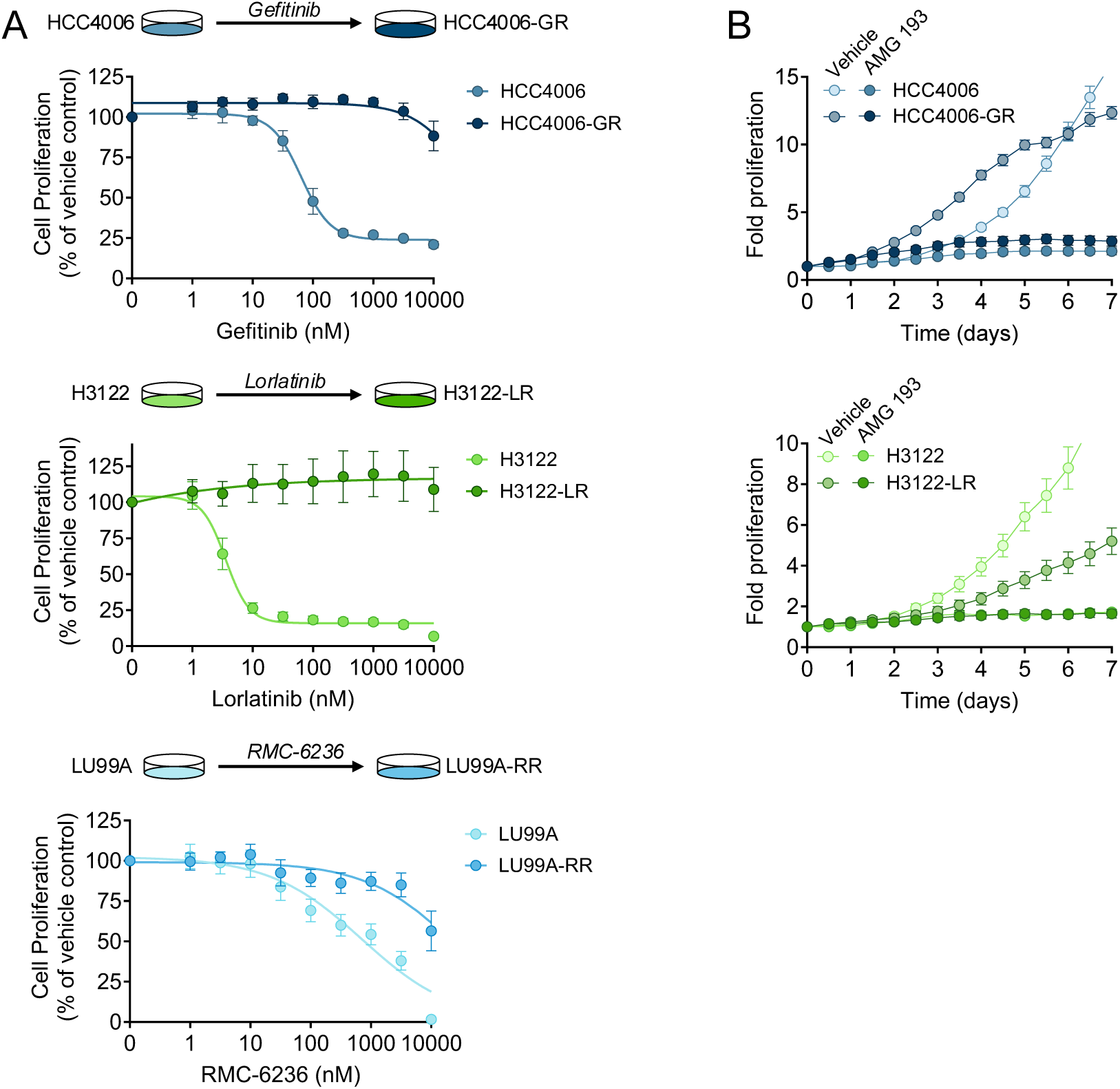
PRMT5 inhibition in drug-tolerant and resistant models. (A) TKI sensitivity in matched parental and resistant NSCLC cell line pairs (HCC4006 vs. HCC4006-GR; H3122 vs. H3122-LR, LU99A vs. LU99A-RR) confirming resistance phenotype. (B) Time-course of proliferation of HCC4006/HCC4006-GR and H3122/H3122-LR cells treated with AMG 193 demonstrating preserved sensitivity across parental and resistant states.

**Supplementary Figure 7.**
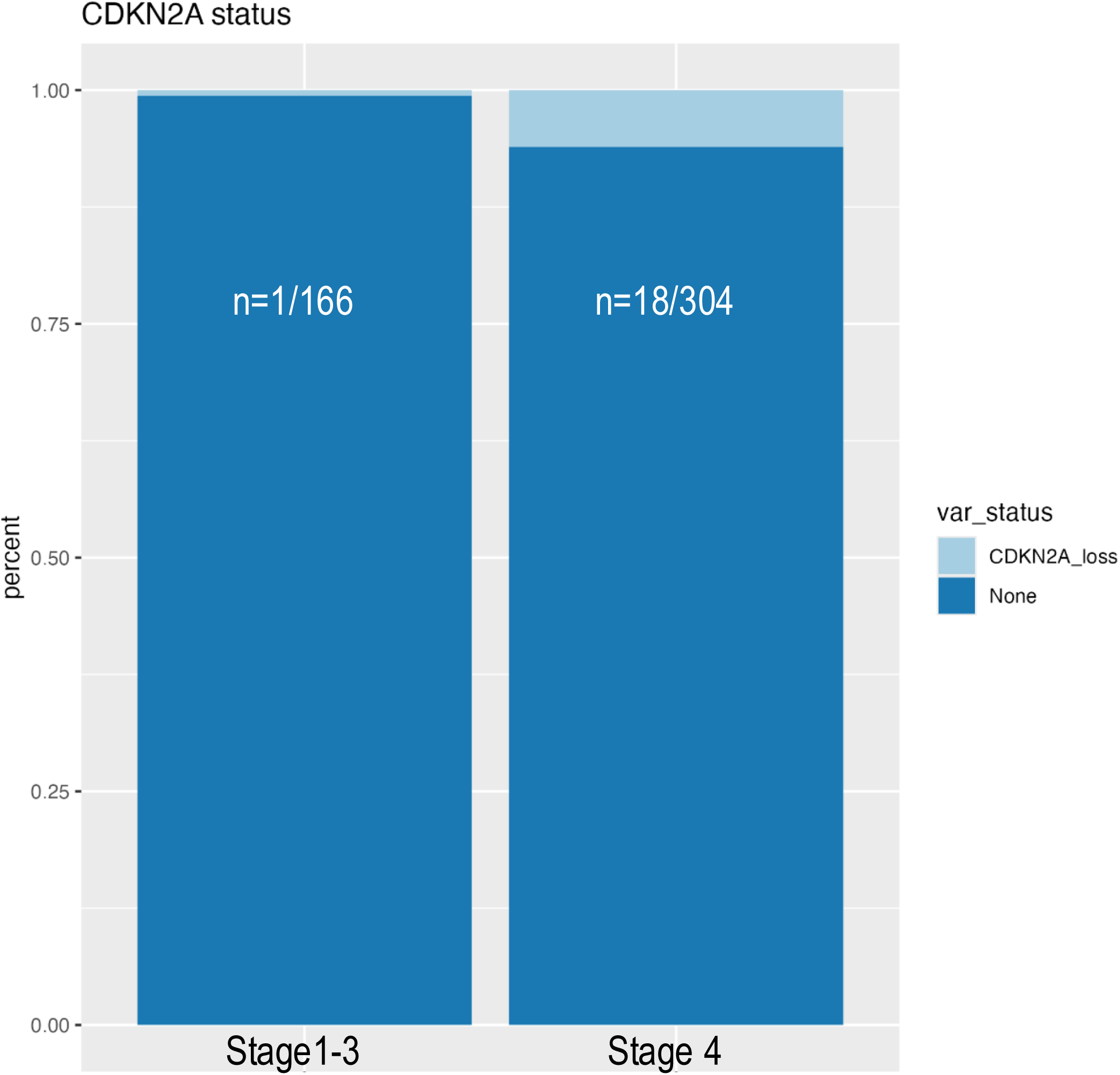
CDKN2A Loss in Early-Stage vs Metastatic NSCLC. Bar graphs depict percentage of lung adenocarcinomas with CDKN2A loss for stage 1-3 tumors vs stage 4 tumors undergoing analysis using Genexus NGS assay. Var_status: Variant (CDKN2A loss) status.

## References

1. Ikushima H, Watanabe K, Shinozaki-Ushiku A, Oda K, Kage H. Pan-cancer clinical and molecular landscape of MTAP deletion in nationwide and international comprehensive genomic data. ESMO Open. Apr 2025;10(4):104535. doi:10.1016/j.esmoop.2025.104535

2. Ashok Kumar P, Graziano SL, Danziger N, et al. Genomic landscape of non-small-cell lung cancer with methylthioadenosine phosphorylase (MTAP) deficiency. Cancer Med. Jan 2023;12(2):1157–1166. doi:10.1002/cam4.4971

3. Brune MM, Roma L, Chijioke O, et al. MTAP expression by immunohistochemistry: a novel biomarker in non-small cell cancer of the lung. J Thorac Oncol. Aug 22 2025;doi:10.1016/j.jtho.2025.08.014

4. Tonini MR. Evaluation of the impact of homozygous MTAP truncations on the activity and selectivity of MTA-cooperative PRMT5 inhibitors. Cancer Research 84(6_Supplement):4631–46312024.

5. Ross JS, Thummalapalli R, Febres-Aldana CA, et al. Clinical Significance of MTAP Deletions and their Overlap with Concurrent Oncogenic Driver Alterations Including EGFR in Non-Small Cell Lung Cancer. J Thorac Oncol. Nov 17 2025;doi:10.1016/j.jtho.2025.11.010

6. Lara-Mejía L, Cardona AF, Mas L, et al. Impact of Concurrent Genomic Alterations on Clinical Outcomes in Patients With ALK-Rearranged NSCLC. J Thorac Oncol. Jan 2024;19(1):119–129. doi:10.1016/j.jtho.2023.08.007

7. Negrao MV, Araujo HA, Lamberti G, et al. Comutations and KRASG12C Inhibitor Efficacy in Advanced NSCLC. Cancer Discov. Jul 07 2023;13(7):1556–1571. doi:10.1158/2159-8290.CD-22-1420

8. Gutiontov SI, Turchan WT, Spurr LF, et al. CDKN2A loss-of-function predicts immunotherapy resistance in non-small cell lung cancer. Sci Rep. Oct 08 2021;11(1):20059. doi:10.1038/s41598-021-99524-1

9. Alessi JV, Wang X, Elkrief A, et al. Impact of Aneuploidy and Chromosome 9p Loss on Tumor Immune Microenvironment and Immune Checkpoint Inhibitor Efficacy in NSCLC. J Thorac Oncol. Nov 2023;18(11):1524–1537. doi:10.1016/j.jtho.2023.05.019

10. Jiang J, Gu Y, Liu J, et al. Coexistence of p16/CDKN2A homozygous deletions and activating EGFR mutations in lung adenocarcinoma patients signifies a poor response to EGFR-TKIs. Lung Cancer. Dec 2016;102:101–107. doi:10.1016/j.lungcan.2016.10.015

11. Guccione E, Richard S. The regulation, functions and clinical relevance of arginine methylation. Nat Rev Mol Cell Biol. 10 2019;20(10):642–657. doi:10.1038/s41580-019-0155-x

12. Kryukov GV, Wilson FH, Ruth JR, et al. MTAP deletion confers enhanced dependency on the PRMT5 arginine methyltransferase in cancer cells. Science. Mar 2016;351(6278):1214–8. doi:10.1126/science.aad5214

13. Mavrakis KJ, McDonald ER, Schlabach MR, et al. Disordered methionine metabolism in MTAP/CDKN2A-deleted cancers leads to dependence on PRMT5. Science. Mar 11 2016;351(6278):1208–13. doi:10.1126/science.aad5944

14. Rodon J, Prenen H, Sacher A, et al. First-in-human study of AMG 193, an MTA-cooperative PRMT5 inhibitor, in patients with MTAP-deleted solid tumors: results from phase I dose exploration. Ann Oncol. Dec 2024;35(12):1138–1147. doi:10.1016/j.annonc.2024.08.2339

15. Engstrom LD, Aranda R, Waters L, et al. MRTX1719 is an MTA-cooperative PRMT5 inhibitor that exhibits synthetic lethality in preclinical models and patients with MTAP deleted cancer. Cancer Discov. Aug 08 2023;doi:10.1158/2159-8290.CD-23-0669

16. Zheng Z, Liebers M, Zhelyazkova B, et al. Anchored multiplex PCR for targeted next-generation sequencing. Nat Med. Dec 2014;20(12):1479–84. doi:10.1038/nm.3729

17. Sheffield BS, Beharry A, Diep J, et al. Point of Care Molecular Testing: Community-Based Rapid Next-Generation Sequencing to Support Cancer Care. Curr Oncol. Feb 23 2022;29(3):1326–1334. doi:10.3390/curroncol29030113

18. Crystal AS, Shaw AT, Sequist LV, et al. Patient-derived models of acquired resistance can identify effective drug combinations for cancer. Science. Dec 19 2014;346(6216):1480–6. doi:10.1126/science.1254721

19. Yoda S, Lin JJ, Lawrence MS, et al. Sequential ALK Inhibitors Can Select for Lorlatinib-Resistant Compound ALK Mutations in ALK-Positive Lung Cancer. Cancer Discov. Jun 2018;8(6):714–729. doi:10.1158/2159-8290.CD-17-1256

20. Zheng S, Wang W, Aldahdooh J, et al. SynergyFinder Plus: Toward Better Interpretation and Annotation of Drug Combination Screening Datasets. Genomics Proteomics Bioinformatics. Jun 2022;20(3):587–596. doi:10.1016/j.gpb.2022.01.004

21. Belmontes B, Slemmons KK, Su C, et al. AMG 193, a Clinical Stage MTA-Cooperative PRMT5 Inhibitor, Drives Antitumor Activity Preclinically and in Patients with MTAP-Deleted Cancers. Cancer Discov. Jan 13 2025;15(1):139–161. doi:10.1158/2159-8290.CD-24-0887

22. Cottrell KM, Briggs KJ, Tsai A, et al. Discovery of TNG462: A Highly Potent and Selective MTA-Cooperative PRMT5 Inhibitor to Target Cancers with MTAP Deletion. J Med Chem. Mar 13 2025;68(5):5097–5119. doi:10.1021/acs.jmedchem.4c03067

23. Marjon K, Cameron MJ, Quang P, et al. MTAP Deletions in Cancer Create Vulnerability to Targeting of the MAT2A/PRMT5/RIOK1 Axis. Cell Rep. Apr 19 2016;15(3):574–587. doi:10.1016/j.celrep.2016.03.043

24. Briggs KJ, Cottrell KM, Tonini MR, et al. TNG908 is a brain-penetrant, MTA-cooperative PRMT5 inhibitor developed for the treatment of MTAP-deleted cancers. Transl Oncol. Feb 2025;52:102264. doi:10.1016/j.tranon.2024.102264

25. Hata AN, Niederst MJ, Archibald HL, et al. Tumor cells can follow distinct evolutionary paths to become resistant to epidermal growth factor receptor inhibition. Nat Med. Mar 2016;22(3):262–9. doi:10.1038/nm.4040

26. Cabanos HF, Hata AN. Emerging Insights into Targeted Therapy-Tolerant Persister Cells in Cancer. Cancers (Basel). May 28 2021;13(11)doi:10.3390/cancers13112666

