## Supplemental Tables for "Prevalence and Actionability of MTAP Loss in Oncogene-Driven Lung Cancer"

### Supplementary Table 1. Baseline Characteristics of Study Population

| Characteristic | Number (%) of Patients<br>Early-Stage Cohort<br>(n = 132) | Number (%) of Patients<br>Metastatic Cohort<br>(n=108) | Number (%) of Patients<br>All Patients<br>(n=240) |
|---|---|---|---|
| **Sex** | | | |
| Male | 46 (35) | 50 (46) | 96 (40) |
| Female | 86 (65) | 58 (54) | 144 (60) |
| **Histology** | | | |
| Adenocarcinoma | 126 (95.5) | 89 (82) | 215 (89.5) |
| Squamous | 1 (1) | 5 (5) | 6 (2.5) |
| Adenosquamous | 2 (1.5) | 3 (3) | 5 (2) |
| Other* | 3 (2) | 11 (10) | 14 (6) |
| **Stage (AJCC v8)** | | | |
| I | 66 (50) | 0 (0) | 66 (50) |
| II | 39 (30) | 0 (0) | 39 (30) |
| III | 27 (20) | 0 (0) | 27 (20) |
| **Molecular Alteration** | | | |
| ALK | 6 (5) | 9 (8) | 15 (6) |
| BRAF | 9 (7) | 3 (3) | 12 (5) |
| EGFR | 43 (33) | 14 (13) | 57 (24) |
| HER2 | 3 (2) | 3 (3) | 6 (3) |
| KRAS | 27 (20) | 30 (28) | 57 (24) |
| MET | 6 (5) | 4 (4) | 10 (4) |
| NRAS | 2 (1) | 0 (0) | 2 (1) |
| NTRK | 1 (1) | 0 (0) | 1 (<1) |
| RET | 3 (2) | 2 (2) | 5 (2) |
| ROS1 | 2 (1) | 6 (6) | 8 (3) |
| None | 30 (23) | 37 (34) | 67 (28) |

*Other: For early-stage: 3 mixed histology (n=2 adenocarcinoma + separate squamous, n=1 adenocarcinoma + large cell neuroendocrine); For metastatic: n=1 large cell neuroendocrine, n=1 sarcomatoid, n=9 non-small cell carcinoma unspecified)

**Supplementary Table 2. Prevalence of MTAP Loss in Lung Cancer Molecular Subsets**

| Molecular Driver Alteration | Early-Stage | Metastatic |
|---|---|---|
| ALK Rearrangement | 0/6 (0%) | 2/9 (22%) |
| BRAF Mutation* | 0/9 (0%) | 1/3 (33%) |
| EGFR Mutation** | 8/43 (18%) | 4/14 (29%) |
| HER2 Alteration | 1/3 (33%) | 1/3 (33%) |
| KRAS Mutation | 3/27 (11%) | 6/30 (20%) |
| MET Alteration | 1/6 (17%) | 1/4 (25%) |
| NRAS Mutation | 0/2 (0%) | --- |
| NTRK Rearrangement | 0/1 (0%) | --- |
| RET Rearrangement | 2/3 (67%) | 0/2 (0%) |
| ROS1 Rearrangement | 0/2 (0%) | 2/6 (33%) |

*BRAF mutation includes class I-III mutations; ***EGFR mutation includes exon 19 deletions, L858R, exon 20 insertions, and atypical mutations; HER2 alterations include mutations and amplification; MET alterations include mutations and amplification

### Supplementary Table 3. Cell Line Characteristics

| Cell Line | Oncogene | Targeted Rx | Resistance mechanism | *CDKN2A* (cBioportal) | *CDKN2A* (Depmap) | *CDKN2A* (MGH NGS) | *CDKN2A* (WB) | *MTAP* (CCLE) | *MTAP* (WB) |
|---|---|---|---|---|---|---|---|---|---|
| PC9 | EGFR exon 19 del | - | Naïve | Diploid | Diploid | | Present | Diploid | Present |
| HCC827 | EGFR exon 19 del | - | Naïve | Diploid | Diploid | | Present | Diploid | Present |
| H3255 | EGFR exon 21 L858R | - | Naïve | Deepdel | Homozygous | | Loss | Deepdel | Loss |
| H1975 | EGFR exon 21 L858R + exon 20 T790M | - | Naïve | Deepdel | Hemizygous | | Loss | Deepdel | Present |
| H1650 | EGFR exon 19 del | - | Naïve | Deepdel | Homozygous | | Loss | Deepdel | Loss |
| HCC4006 | EGFR exon 19 del | - | Naïve | Deepdel | Homozygous | | Loss | Deepdel | Loss |
| MGH119-1 | EGFR exon 19 del | - | Naïve | | | Unknown | Loss | | Loss |
| MGH121-1 | EGFR exon 19 del | Erlotinib | Exon 20 T790M | | | Unknown | Present | | Present |
| MGH707-1 | EGFR exon 19 del | Erlotinib, Afatinib | Exon 20 T790M | | | Unknown | Loss | | Loss |
| MGH1157-1 | EGFR exon 19 del | - | Naïve | | | Loss | Loss | | Present |
| MGH1157-3 | EGFR exon 19 del | Nazartinib + Gefitinib | MET Amplification | | | Loss | Loss | | Present |
| MGH1167-1 | EGFR exon 18 G719A + exon 20 R776S | - | Naïve | | | Loss | Loss | | Present |
| MGH10063-2 | EGFR exon 19 del | Gefitinib, Osimertinib, Osimertinib + Savolitinib | Exon 20 T790M | | | Loss | Loss | | Present |
| MGH10077-3 | EGFR exon 19 del | Erlotinib, Osimertinib | Exon 20 T790M, C797S | | | Loss | Loss | | Loss |
| MGH10016-4 | EGFR exon 21 L858R | Nazartinib | Not identified | | | Loss | Loss | | Loss |
| MGH10016-5 | EGFR exon 21 L858R | Nazartinib, Osimertinib | MET amplification | | | Loss | Loss | | Loss |
| MGH794-5 | EGFR exon 19 del | Erlotinib, Nazartinib, Osimertinib | Exon 20 T790M | | | Loss | Loss | | Loss |
| MGH10155-1 | EGFR exon 19 del | - | Naïve | | | Loss | Loss | | Present |
| MGH10155-6 | EGFR exon 19 del | Osimertinib | Not identified | | | Loss | Loss | | Present |
| H23 | KRAS G12C | - | Naïve | Deepdel | Hemizygous | | Loss | Unknown | Present |
| H2030 | KRAS G12C | - | Naïve | Deepdel | Hemizygous | | Loss | Deepdel | Present |
| H2122 | KRAS G12C | - | Naïve | Deepdel | Homozygous | | Loss | Unknown | Present |
| LU65 | KRAS G12C | - | Naïve | Diploid | Diploid | | Present | Unknown | Present |
| H1792 | KRAS G12C | - | Naïve | Deepdel | Hemizygous | | Present | Deepdel | Present |
| LU99A | KRAS G12C | - | Naïve | Deepdel | Homozygous | | Loss | Deepdel | Loss |
| SW1573 | KRAS G12C | - | Naïve | Deepdel | - | | Loss | Deepdel | Loss |
| CALU1 | KRAS G12C | - | Naïve | Diploid | Diploid | | Loss | Diploid | Present |
| H358 | KRAS G12C | - | Naïve | Diploid | - | | Loss | Diploid | Present |
| H1573 | KRAS G12A | - | Naïve | Diploid | - | | Loss | Diploid | Present |
| H2009 | KRAS G12A | - | Naïve | Diploid | - | | Present | Diploid | Present |
| SKLU1 | KRAS G12D | - | Naïve | Deepdel | - | | Loss | Deepdel | Loss |
| A427 | KRAS G12D | - | Naïve | Deepdel | - | | Loss | Unknown | Present |
| A549 | KRAS G12S | - | Naïve | Deepdel | Homozygous | | Loss | Deepdel | Loss |
| H441 | KRAS G12V | - | Naïve | Deepdel | | | Loss | Deepdel | Present |
| SW900 | KRAS G12V | - | Naïve | Deepdel | Homozygous | | Loss | Deepdel | Present |
| COR-L23 | KRAS G12V | - | Naïve | Deepdel | Diploid | | Loss | Deepdel | Present |
| H1944 | KRAS G13D | - | Naïve | Deepdel | - | | Loss | Deepdel | Present |
| DV90 | KRAS G13D | - | Naïve | Diploid | - | | Loss | Diploid | Present |
| H1155 | KRAS Q61H | - | Naïve | Diploid | - | | Present | Diploid | Present |
| H460 | KRAS Q61H | - | Naïve | Deepdel | Homozygous | | Loss | Diploid | Present |
| MGH1070-1 | KRAS G12S | - | Naïve | | | Unknown | Present | | Present |
| MGH1088-1 | KRAS G12C | - | Naïve | | | Unknown | Loss | | Present |
| MGH1089-1 | KRAS G12D | - | Naïve | | | Unknown | Loss | | Loss |
| MGH1112-1 | KRAS G12C | - | Naïve | | | Unknown | Present | | Present |
| MGH1114-1 | KRAS G12C | - | Naïve | | | Unknown | Present | | Present |
| MGH1138-1 | KRAS G12C | - | Naïve | | | Unknown | Loss | | Loss |
| MGH1143-2 | KRAS G12C | - | Naïve | | | Unknown | Loss | | Present |
| MGH9029-1 | KRAS G12C | - | Naïve | | | Unknown | Loss | | Loss |

### Supplementary Table 3. Cell Line Characteristics (Continued)

| | | | | | | | | | *MTAP* (WB) |
|---|---|---|---|---|---|---|---|---|---|
| H3122 | EML4-ALK v1 | - | Naïve | Deepdel | Homozygous | | Loss | Unknown | Loss |
| MGH006-1 | EML4-ALK v1 | - | Naïve | | | Unknown | Present | | Present |
| MGH026-1 | EML4-ALK v3 | - | Naïve | | | Unknown | Loss | | Present |
| MGH048-1 | EML4-ALK v1 | - | Naïve | | | Unknown | Present | | Present |
| MGH064-1 | EML4-ALK v2 | - | Naïve | | | Unknown | Loss | | Present |
| MGH065-1 | EML4-ALK v1 | Crizotinib | FGFR1 | | | Unknown | Loss | | Present |
| MGH073-2 | EML4-ALK v3 | Crizotinib | FGFR1 | | | Unknown | Loss | | Loss |
| MGH075-2 | EML4-ALK v2 | Ceritinib | Not identified | | | Unknown | Loss | | Present |
| MGH079-2 | EML4-ALK v1 | Crizotinib, Alectinib | RTK bypass | | | Unknown | Loss | | Loss |
| MGH902-1 | EML4-ALK v3 | Crizotinib, Ceritinib | MDR1 overexpression | | | Unknown | Present | | Present |
| MGH915-3 | EML4-ALK v1 | Ceritinib, Alectinib | Not identified | | | Loss | Loss | | Loss |
| MGH947-1 | EML4-ALK v5' (E18; A20) | Crizotinib, Brigatinib, Alectinib, Lorlatinib | Not identified | | | Unknown | Loss | | Present |
| MGH952-1 | EML4-ALK v5' (E18; A20) | Brigatinib | MET overexpression | | | Unknown | Loss | | Present |
| MGH953-1 | EML4-ALK v3 | Crizotinib | Not identified | | | Unknown | Loss | | Present |
| MGH979-6.2 | EML4-ALK v3 | Alectinib, Ceritinib | HGF overexpression | | | Unknown | Loss | | Loss |
| MGH990-2 | EML4-ALK v3 | Crizotinib, Alectinib, Lorlatinib | Not identified | | | Loss | Loss | | Loss |
| MGH9092-1 | EML4-ALK v5' (E18; A20) | Crizotinib, Alectinib, Lorlatinib | Not identified | | | Unknown | Loss | | Present |
| MGH9162-1 | EML4-ALK v3 | Alectinib | Not identified | | | Loss | Loss | | Present |
| MGH9243-1 | EML4-ALK v1 | Alectinib, Lorlatinib | HGF overexpression | | | Unknown | Loss | | Present |
| MGH10029-1 | ALK Rearrangement (other) | - | Naïve | | | Loss | Present | | Present |
| MGH10371-1 | EML4-ALK v1 | Alectinib, Lorlatinib | Not identified | | | Loss | Loss | | Present |
| MGH9018-1 | ROS1 Rearrangement | Crizotinib | ROS1 G2032R | | | Loss | Loss | | Present |
| MGH10366-2 | ROS1 Rearrangement | Crizotinib, lorlatinib, Zidesamtinib | Not identified | | | Loss | Present | | Present |
| MGH9174-1 | KIF5B-RET Rearrangement | Selpercatinib | RET G810S | | | Loss | Loss | | Present |
